# Redundant and specific roles of cohesin STAG subunits in chromatin looping and transcription control

**DOI:** 10.1101/642959

**Authors:** Valentina Casa, Macarena Moronta Gines, Eduardo Gade Gusmao, Johann A. Slotman, Anne Zirkel, Natasa Josipovic, Edwin Oole, Wilfred F.J. van IJcken, Adriaan B. Houtsmuller, Argyris Papantonis, Kerstin S. Wendt

## Abstract

Cohesin is a ring-shaped multiprotein complex that is crucial for 3D genome organization and transcriptional regulation during differentiation and development. It also confers sister chromatid cohesion and facilitates DNA damage repair. Besides its core subunits SMC3, SMC1A and RAD21, cohesin contains in somatic cells one of two orthologous STAG subunits, SA1 or SA2. How these variable subunits affect the function of the cohesin complex is still unclear. SA1- and SA2-cohesin were initially proposed to organize cohesion at telomeres and centromeres, respectively. Here, we uncover redundant and specific roles of SA1 and SA2 in gene regulation and chromatin looping using HCT116 cells with an auxin-inducible degron (AID) tag fused to either SA1 or SA2. Following rapid depletion of either subunit, we perform high resolution Hi-C, RNA-sequencing and sequential ChIP studies to show that SA1 and SA2 do not co-occupy individual binding sites and have distinct ways how they affect looping and gene expression. These findings are supported at the single cell level by single-molecule localizations via dSTORM super-resolution imaging. Since somatic and congenital mutations of the SA subunits are associated with cancer (SA2) and intellectual disability syndromes with congenital abnormalities (SA1 and SA2), we verified SA1-/SA2-dependencies using human neural stem cells, hence highlighting their importance for understanding particular disease contexts.

## Introduction

Cohesin is a multiprotein complex with fundamental roles in genome stability and in spatial organization of the chromatin fiber within the cell nucleus. The ring shape of the complex is determined by its core subunits SMC3, SMC1A, RAD21 (Smc3, Smc1 and Scc1 in *S. cerevisiae*) and by one SA/STAG (Stromal Antigen) subunit (Scc3 in *S. cerevisiae*). Furthermore, transiently interacting subunits and regulators of this ring-like complex are PDS5A/PDS5B (Pds5 in *S. cerevisiae*), Sororin, WAPL and Kollerin, consisting of NIPBL and MAU2 (Ciosk et al. 2000).

Vertebrates have three SA/STAG subunit orthologues, SA1 (also STAG1), SA2(also STAG2) and SA3 (also STAG3), with SA3 only expressed in germ cells and certain cancers (Prieto et al. 2001; Winters et al. 2014). SA1 and SA2 are ubiquitously expressed in somatic cells at ratios that may vary among different cell types and are mutually exclusively incorporated in the complex (Losada et al. 2000). SA1 and SA2 are overall highly similar at the protein sequence level (∼75% amino acids conserved) with only their N- and C-termini diverging (30-50% homology) (Losada et al. 2000; Kong et al. 2014).

Previously, SA1 and SA2 have been reported to be differentially required for sister chromatid cohesion at telomeres and centromeres, respectively (Canudas and Smith 2009), while both contribute to chromosome arm cohesion (Remeseiro et al. 2012). STAG1-null mouse embryonic fibroblasts show increased aneuploidy as a result of telomeric cohesion defects (Remeseiro et al. 2012). In addition, SA1 protects chromosome ends and promotes replication by facilitating restart from stalled replication forks (Canudas and Smith 2009; Deng et al. 2012; Remeseiro et al. 2012). On the other hand, acute SA2 depletion results in partial loss of centromeric cohesion (Canudas and Smith 2009; Remeseiro et al. 2012; Lennox and Behlke 2016; Busslinger et al. 2017). Moreover, SA2 is required for DNA damage repair via the homologous recombination pathway in the S/G2 cell cycle phases (Kong et al. 2014), but both SA1 and SA2 are necessary for the S-phase DNA damage checkpoint (Kong et al. 2014). Cells can tolerate the absence of either SA1 or SA2 alone but not of both simultaneously, suggesting that SA1 and SA2 have at least partially redundant roles (Hill et al. 2016; Benedetti et al. 2017; van der Lelij et al. 2017).

Besides their apparent functional differences, cohesin complexes containing SA1 or SA2 colocalize with the CTCF chromatin insulator throughout the mouse and human genomes (Wendt et al. 2008) (Rubio et al. 2008; Remeseiro et al. 2012). Cohesin and CTCF form chromatin loops by tethering distant genomic regions (Hadjur et al. 2009; Nativio et al. 2009; Zuin et al. 2014a; Nora et al. 2017; Rao et al. 2017) preferentially between cohesin/CTCF sites with convergently orientated CTCF DNA motifs (Rao et al. 2014). Further partitioning of the genome into loop clusters, known as topologically-associating domains (TADs) (Dixon et al. 2012; Nora et al. 2012), impacts gene regulation by facilitating or abrogating promoter-enhancer contacts. How SA1-cohesin and SA2-cohesin are involved in shaping this spatial organization of the genome remains unclear. Decoding this is will not only complete our understanding of cohesin molecular functions, but will allow us to better understand different pathologies linked to these orthologues. Amongst all cohesin subunits and co-factors, SA2 is the one most frequently mutated in different cancer types, such as urothelial bladder carcinoma (26% of tumor samples) (Balbas-Martinez et al. 2013; Solomon et al. 2013), Ewing sarcoma (15-22% of tumor samples) (Brohl et al. 2014; Tirode et al. 2014) and or acute myeloid leukemia (6% of tumor samples) (Rocquain et al. 2010). Interestingly, congenital loss-of-function mutations in SA1 and SA2 do not cause Cornelia de Lange syndrome (CdLS), the major syndrome linked to mutations in core cohesin components and co-regulators, but rather associate with developmental delay and various malformations (Lehalle et al. 2017; Mullegama et al. 2017; Soardi et al. 2017).

Here, we combined chromatin genomics, in Hi-C and ChIP-seq, with single-molecule approaches, in dSTORM imaging, to decipher the individual contributions of SA1 and SA2 in the 3D organization of the genome at different scales.

## Results

### Acute auxin-inducible degradation of SA1 and SA2 in human diploid cells

In order to generate cell lines that allow for the rapid depletion of either SA1 or SA2, but also for their visualization and immunopurification, we inserted the mini auxin-inducible degron (AID) tag in frame with mClover at the 3’ends of the *SA1* and *SA2* open reading frames in HCT-116-CMV-OsTIR1 cells (Natsume et al. 2016b) (**Fig. 1A,B**). These cell lines, constitutively-expressing the TIR1 ubiquitin ligase, are hereafter referred to as SA1-AID and SA2-AID. Thus, induction of acute proteasomal degradation of either SA1 or SA2 fused to the mini-AID tag via ubiquitination is triggered by addition of the plant hormone, auxin. Immunoprecipitation experiments showed that the AID-mClover-tagged SA1 and SA2 subunits are properly incorporated in the cohesin complex (**Fig. 1C, D**) and that both are quantitatively degraded upon addition of auxin, without reciprocally affecting the levels of the non-targeted STAG subunit or those of SMC1A (**Fig. 1E-H**). In all experiments with SA1-AID and SA2-AID cells we used an auxin treatment for 12 hours. To show that both SA1 and SA2 were virtually fully removed from chromatin in response to auxin we used ChIP-qPCR (**Fig. 1I, J**). Thus, we established two HCT116-based cell lines, where SA1 or SA2 can be essentially fully depleted from chromatin.

**Figure 1.**
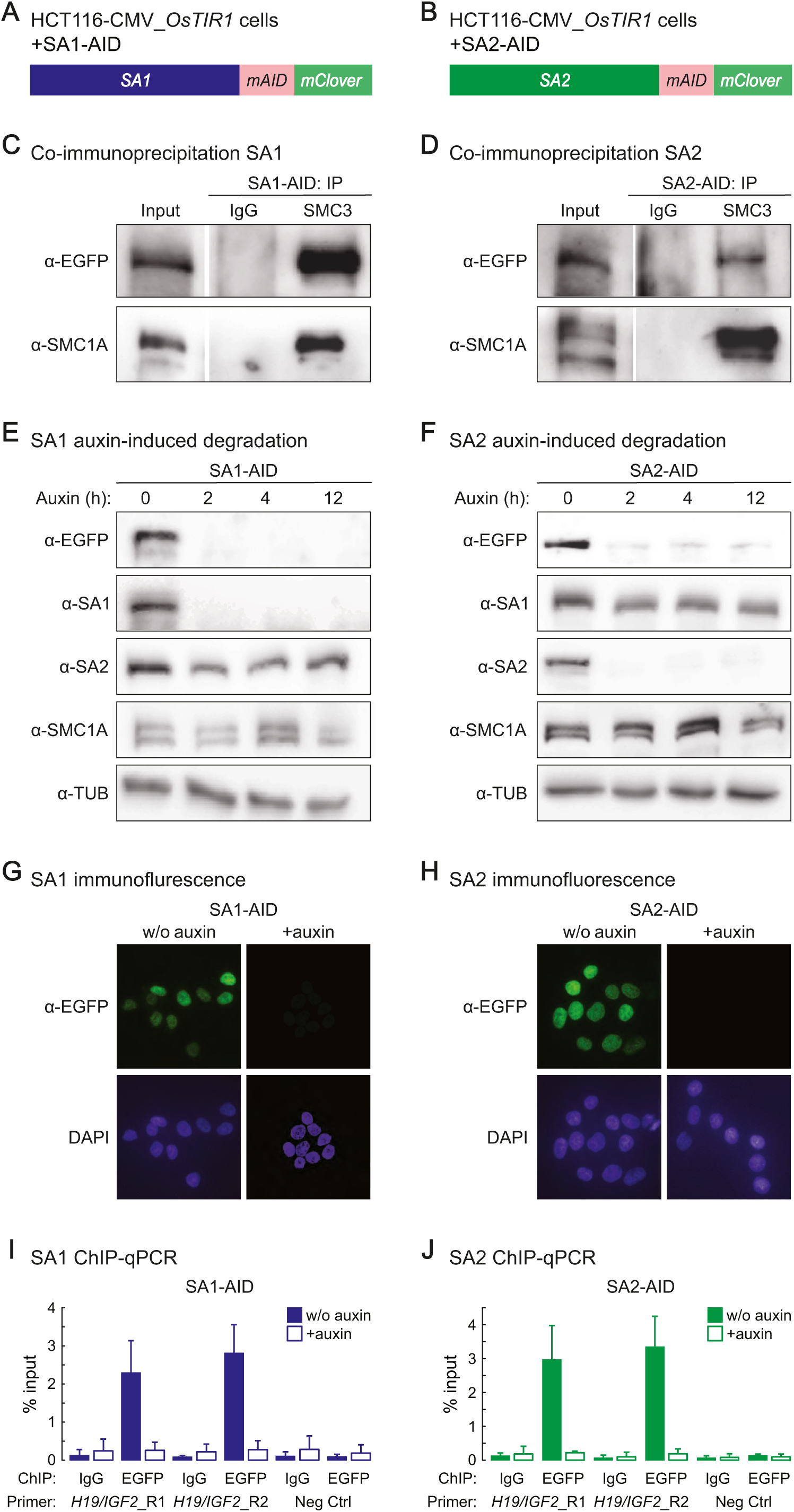
Acute auxin-inducible degradation of SA1 and SA2. **(A)** Schematic of the CRIPR-mediated modification of SA1 and SA2 to add a mini auxin-inducible degron tag (mAID) and a mClover tag to the C-termini of SA1 and SA2 in HCT116-CMV-OsTIR1 cells. **(B)** Immunoprecipitation of SMC3 from SA1-AID and SA2-AID cells demonstrates that the modified subunits are incorporated in the cohesin complex. **C**) Addition of auxin leads to a degradation of SA1-AID and SA2-AID within 2 hours without affecting the protein level of the respective other ortholog. **D**) Visualization of the SA1-AID and SA2-AID degradation 2 hours after auxin addition by immunostaining. **E**) ChIP-qPCR with anti-EGFP antibodies (detects mClover) shows efficient depletion of SA1-AID and SA2-AID from cohesin sites in the H19/IGF2 locus.

### SA1 and SA2 display both overlapping and discrete binding positions on chromatin

To obtain a comprehensive repertoire of SA1 and SA2 binding on chromatin, we performed ChIP-seq using antibodies against EGFP that also detect the EGFP-derivative mClover fused to each SA subunit (Natsume et al. 2016a). Following data analysis, we identified 17,615 and 13,524 robust binding sites for SA1 and SA2, respectively (q-value cutoff <0.005; **Fig. 2A**). More than 70% of the recorded binding sites showed ChIP-seq signal for both SA1 and SA2 (SA1/2), whereas exclusive SA1 (SA1-only) and SA2 binding (SA2-only) was observed for 4,650 and 568 sites, respectively (**Fig. 2B**). Nonetheless, all positions expectedly aligned to cohesin-bound (RAD21) sites, consistent with the 1:1:1:1 stoichiometry of subunits in cohesin complexes, as well as with CTCF-bound sites (from HCT116-RAD21-mAC cells; (Natsume et al. 2016b; Rao et al. 2017) (**Figs. 2B** and **S1A**). The overall signal intensity of CTCF at SA1/SA2 common peaks was higher than in SA1-only and SA2-only peaks, which was also true for the respective SA1 and SA2 signals, possibly indicative of lower occupancy at this subset of sites (**Fig. 2C**). We also interrogated exemplary loci using ChIP-qPCR to confirm subunit-specific binding of SA1 and SA2 at SA1-/SA2-only sites. We found that binding was abolished upon auxin treatment only in the relevant mAID line (**Figs. 2D,E** and **S1B-F**), and that recruitment of cohesin core subunits, as exemplified by SMC3, depended on the presence of the respective auxin-sensitive STAG subunit at SA1-/SA2-only sites (**Fig. S1D,F**).

**Figure 2.**
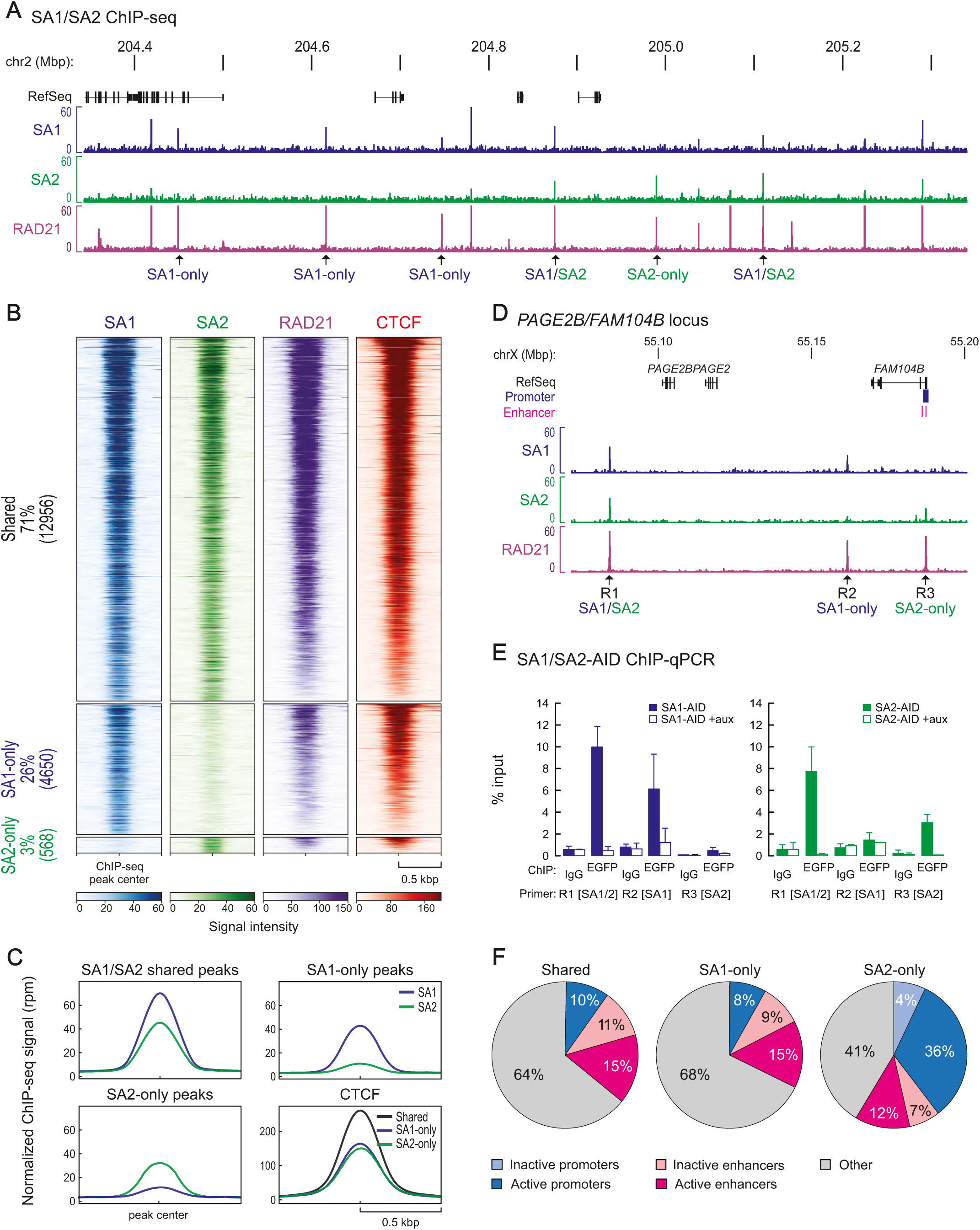
SA1 and SA2 have overlapping as well as discrete binding positions on chromatin. **A)** ChIP-sequencing profiles of SA1 and SA2 from SA1-AID and SA2-AID cells using anti-EGFP antibodies are displayed together with published RAD21 ChIP-seq data (Rao et al. 2017). Examples of SA1-only, SA2-only and SA1/2 shared sites are indicated. **B**) Heatmaps showing the ChIP-seq signals for SA1, SA2, Rad21 and CFCF in common and only sites. RAD21 and CTCF ChIP-seq data are from Rao et al., 2017 (Rao et al. 2017). **C**) Averaged signal intensity of SA1, SA2 and CTCF on SA1/2 common sites and SA1/SA2-only sites. **D**) Examples of SA1 and SA2 binding sites in the PAGE2B/FAM104B locus that were tested in (E) by ChIP-qPCR. **E**) Efficiency of auxin-mediated degradation was tested by ChIP-qPCR for different SA1/SA2 sites. **F**) Localization of SA1/SA2 common sites or SA1 or SA2-only sites to inactive/active promoters or active/inactive enhancers based on the presence of characteristic histone modifications (Table S7). The percentage of the sites in the different regions is indicated.

We next asked whether either STAG subunit was replaced by the other after auxin-induced degradation. SA1 ChIP-qPCR in SA2-AID cells showed such reciprocal replacement at every SA1/2 site with an average 1.5-fold increase after SA2 degradation. On the other hand, SA2-only sites showed less, if any, enrichment of SA1 (**Fig. S1G, H**). The converse experiment, where SA2 ChIP-qPCR was performed after SA1 degradation, showed that SA2 may only minimally, if at all, substitute for SA1 (**Fig. S1I, J**). To address this discrepancy between SA1-only and SA2-only sites, we looked into their distributions across the genome, focusing on regulatory regions. Almost 10% of the SA1/2 common sites are found at active gene promoters and ∼26% at enhancers (15 % at active enhancers). For SA1-only sites we observed a similar distribution, 8% at active promoters and 24 % at enhancers (15% active enhances). For SA2-only sites we observe a significantly larger fraction of 33% at active promoters and 19% at enhancers (12% active enhancers) (*P*<0.001, Fisher’s exact test; **Fig. 2F** and **Table S7**). Analysis of transcription factor motifs contained in DNase I-hypersensitive footprints under all SA1/SA2 ChIP-seq peaks showed that they potentially co-bind with a diverse and non-overlapping repertoire of transcription factors (**Fig. S2A**). A similar analysis of SA1-/SA2-only peaks recapitulated these divergent repertoires (**Fig. S2D**). A GO-term analysis for the different transcription factor found here using Metascape (Zhou et al. 2019) revealed that these are implicated in development and differentiation processes. Using published ChIP-seq data from HCT116 cells (see Table S8), we could show that SA2 peaks, compared to SA1 peaks, are more associated with H3K4me3 signal, indicating active promoters, and less with H3K4me1, indicative for enhancers (**Fig. S2B**). This is consistent with the different preferential localizations of the SA1- or SA2-only sites (**Fig. 2F**). We could also confirm the observation that colocalizing transcription factors have differential preferences for SA1- and SA2- only sites, eg. FOSL1 (**Fig. S2C**) confirms the observation that FOS/JUN family proteins have a preference for SA1-only peaks (**Figs. S2A and D**).

### SA1 and SA2 do not co-occupy individual binding sites

Given that >70% of all SA1 and SA2 ChIP-seq peaks show binding by both STAG subunits, we sought to address whether cohesin complexes containing SA1 and SA2 indeed localize together at SA1/2 binding sites, or whether signal colocalization reflects the average of non-overlapping binding at individual alleles across the cell population used in ChIP. We performed sequential ChIP (“Re-ChIP”) experiments using an optimized protocol. We first performed a chromatin immunoprecipitation from HCT116-SA1-AID cells using antibodies against SA2, EGFP (mClover), SMC3 and rabbit IgG that were crosslinked to beads to avoid carry-over into the next reaction. Then, a second ChIP reaction takes place using antibodies directed against SA1 or SA2 (**Fig. 3A**). Using this approach, we found that we essentially pull-down no SA1-bound chromatin from a pool of complexes binding SA2, and *vice versa* barely any SA2-bound chromatin from a pool of SA1 complexes (**Fig. 3B, C**). Still, in a control experiment, both SA1- and SA2-bound chromatin can be immunoprecipitated from an eluate of an anti-SMC3 ChIP (**Fig. S3A, B**). This shows that SA1-cohesin and SA2-cohesin rings do not co-occupy the same positions on the individual DNA alleles.

**Figure 3.**
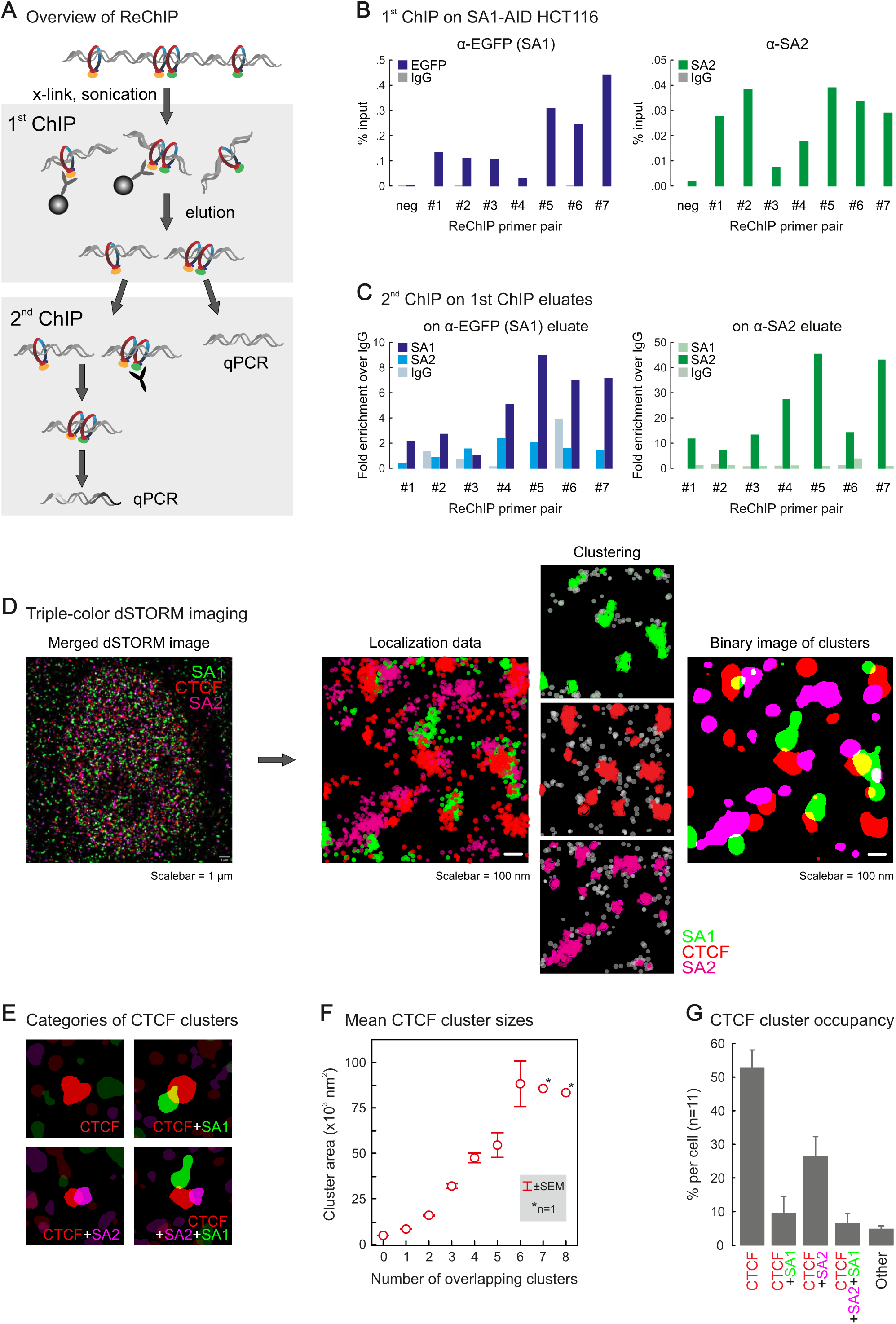
SA1 and SA2 do not co-occupy individual binding sites. **A)** Overview of the re-ChIP protocol. Please note that in the 1^st^ ChIP the antibodies were crosslinked to the beads, allowing for their complete removal from the eluate. **B)** First ChIP in the Re-ChIP protocol performed from SA1-AID cells with antibodies against EGFP to precipitate SA1 and against SA2. For the analysis, qPCR primers directed against one negative and seven positive cohesin binding sites were used. **C)** The 2^nd^ CHIP performed was performed with control IgG, anti-SA1 and anti-SA2 antibodies using the eluates from the 1^st^ CHIP. The same primers as in (B) were used for the analysis. **D)** Analysis pipeline of the dSTORM imaging data. A merged reconstructed triple color dSTORM image of a single nucleus stained for SA1 (green), SA2 (red) and CTCF (magenta) is shown at the left side. For a zoomed region from the same cell the raw localizations are shown as scatter plot. The result of the kernel density estimation-based clustering algorithm for the individual colors are shown next to it. Not clustered localizations are depicted in grey. Finally, a binary image of the same region is shown at the right with the clusters for all three proteins represented. **E)** Examples of the four most common groups of CTCF cluster with either: none, one SA1, one SA2, or both one SA1 and one SA2 cluster adhering. **F)** The cluster area size (nm^2^) for CTCF is plotted grouped per number of adherent clusters of SA1 or SA2. **G)** Frequency of CTCF cluster groups per cell measured for 11 cells. All CTCF clusters falling in other groups are depicted as “Others”. This group contains 18 subgroups, eg. adhering to more than 2 clusters or 2 clusters of the same SA. All error bars show standard error of the mean (+/- SEM).

We followed this up by single molecule localization studies using triple-color dSTORM (direct Stochastical Optical Reconstruction Microscopy; (van de Linde et al. 2011)), a method that allows for the localization of single molecules with approximately 25 nm precision. We performed triple immunostaining for CTCF, SA1 and SA2 in human primary skin fibroblasts, ideal for imaging due to their flat morphology. Localization data were analyzed and clustered (Paul et al. 2019), and the binary images generated were used to infer CTCF, SA1 and SA2 co-clustering (**Fig. 3D**). In this analysis, we focused on correlations relative to the CTCF signal, as nearly all CTCF molecules are chromatin-bound (**Fig. S3E**) and the majority of cohesin sites are also occupied by CTCF (**Fig. 2B**). In our clustered data, we observed CTCF clusters that either stand alone or overlap those by SA1 or SA2 (**Fig. 3E**) and display a mean size of 12,100 nm^2^. Standalone CTCF clusters consistently show a size of ∼5,300 nm^2^ (the smallest observed) and represent more than half of all observed CTCF clusters (**Fig. 3F, G**). Since CTCF-CTCF loops are typically stabilized by at least one cohesin complex, we postulated that such standalone clusters could represent localization signals from single CTCF molecules. This premise was substantiated by CTCF dSTORM imaging in HCT116-RAD21-mAC cells, where auxin-induced RAD21 degradation leads to a complete loss of CTCF-CTCF loops genome-wide (Rao et al. 2017). Comparison of CTCF cluster sizes in these cells in the absence or presence of auxin, showed a reduction in the mean CTCF cluster size from ∼11,500 to 6,400 nm^2^ (**Fig. S3C, D**), similar to what was observed for all and for standalone CTCF clusters in fibroblasts, respectively (**Fig. 3F**).

CTCF clusters overlapping adjacent STAG clusters appear progressively larger as more SA1 and/or SA2 associate with them (ranging from ∼8,800 for overlap with a single cluster to >50,000 nm^2^ for overlap with 5 SA1/SA2 clusters; **Fig. 3F**), and this is most likely indicative of higher-order chromatin structures harboring multiple proteins. However, and in agreement with our Re-ChIP data, ∼26% CTCF clusters overlap SA2 and ∼10% overlap SA1, compared to the ∼6% overlapping both SA1 and SA2 (**Fig. 3G**). Taken together, these analyses highlight the relative dynamics of SA1- and SA2-containing cohesin complexes and, to the extent that we record higher-order chromatin interactions, these seem to predominantly involve only one of the two STAG proteins.

### SA1 and SA2 differentially contribute to higher-order chromatin organization

It was previously shown that auxin-induced cohesin degradation completely alleviates CTCF-CTCF loop formation genome-wide leading to the profound rearrangement of regulatory chromatin interactions (Rao et al. 2017). Here, we generated highly correlated *in situ* Hi-C data from two independently-obtained HCT116-SA1/SA2-AID lines (**Fig. S4A**) that afford far higher resolution than previously reported Hi-C from STAG-knockdown cells (Kojic et al. 2018), allowing us to now assess the individual contribution of SA1 and SA2 in spatial chromatin organization at different scales.

First, we subtracted Hi-C contact matrices derived from auxin-treated and untreated cells at a 200-kbp resolution in a binarized fashion (see Methods for details); this revealed that changes induced to higher-order chromosomal organization by SA1 or SA2 depletion are not only non-converging, but at places display inverse patterns (see an examples from chr17 in **Fig. 4A,B** and a zoom-in in chr13 **Fig. 4C, D**). However, we detected no notable switches between A- and B-compartments upon either depletion, consistent with previous observations by us and others upon cohesin or CTCF loss (Seitan et al. 2013; Sofueva et al. 2013; Zuin et al. 2014a; Nora et al. 2017; Rao et al. 2017) (**Fig. S4B**). Still, a general increase in longer-range interactions (>2Mb) was seen in SA1- depleted cells, in contrast to a largely inversed trend in SA2-depleted ones (**Fig. S4C**).

**Figure 4.**
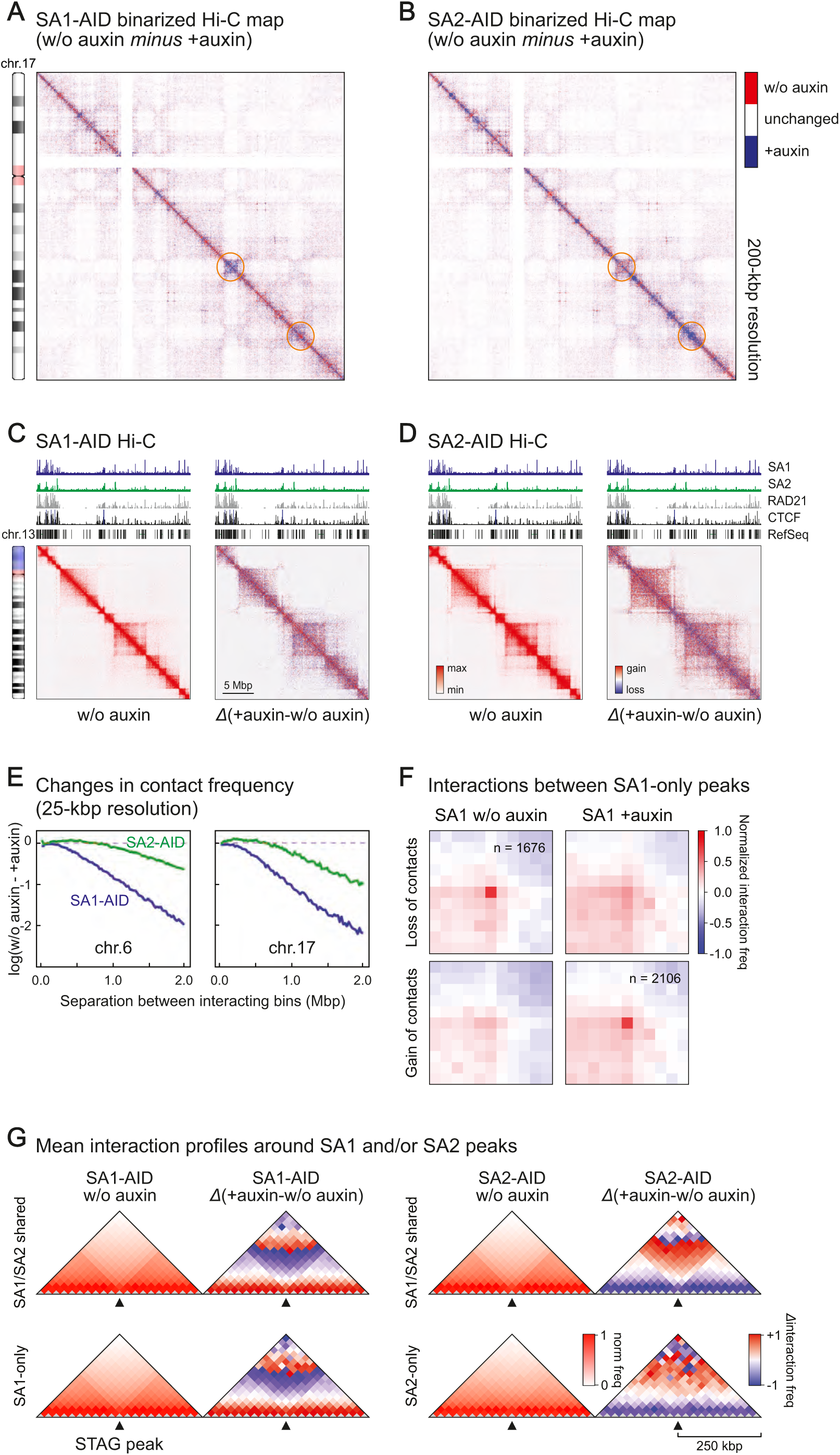
Hi-C reveals different contributions of SA1 and SA2 to higher order chromatin structure. **A, B)** Binarized maps of chromosome 17 illustrating the chromatin contact changes after auxin-mediated degradation of SA1 (**A,** SA1-AID) or SA2 (**B,** SA2-AID). Areas with particularly striking changes are encircled. **C, D)** Zoom-in for a region of chromosome 13 into the contact maps of untreated SA1-AID and SA2-AID cells and the corresponding differential interaction maps after auxin-mediated degradation. **E)** Change of chromatin contact frequency relative to the separation of the contacting bins (25 kb) after SA1 or SA2 degradation for the examples of chromosomes 6 and 17. **F)** Aggregate peak analysis of SA1-only peaks showing that a lot of the sites lose their respective contacts after SA1 degradations while alternative contacts are formed. **G)** Insulation plots showing the averaged Hi-C signals for all SA1/SA2 shared peaks as well as all SA1-only and SA2-only peaks in the untreated maps (w/o auxin). For the same peaks the averaged differential HiC signals in the different maps are shown. The maps are presented with a bin size of 25k and 10 bins to the left and right of each binding site.

Looking at the level of individual TADs (up to ∼2 Mbp), SA1 depletion resulted in a more dramatic loss of interactions and a general contact reshuffling compared to SA2 depletion (**Figs. 4E** and **S4D**). Using TADtool (Kruse et al. 2016) to identify TAD boundaries, we found that the majority (∼60%) remained unaffected despite SA1 or SA2 depletion. Of the ∼40% TAD boundaries that did change, most showed merging with an adjacent TAD (i.e., loss of local insulation in ∼20% TADs in either SA1- or SA2-depleted cells) or the appearance of a new boundary (in ∼15% TADs in either case; **Fig. S4E**), an effect consistent with the collapse of loop domains seen upon RAD21 depletion (Rao et al. 2017).

We finally used our Hi-C data to plot average interaction profiles around SA1/SA2-shared peaks and SA1- or SA2-only peaks (**Fig. 4G**). In non-auxin-treated cells SA1/SA2-shared peaks and SA1-only, but not SA2-only sites, appear to reside at sites of local contact insulation (i.e., at TAD or loop domain boundaries). Then, upon depletion of either subunit, we again observe shorter-range contact changes around SA2-only sites and longer-range changes around SA1-only and SA1/SA2-shared sites (**Fig. 4G**).

We next used the highest (i.e., 10-kbp) resolution our Hi-C data could afford to directly query looping interactions forming between sites bound by one or the other cohesin-STAG subcomplex. Because of the high redundancy of SA1- and SA2-binding to chromatin, we focused on SA1-only and SA2-only peaks, where effects from auxin-induced degradation of each subunit would be accentuated. Surprisingly, and with the caveat of their low numbers (568 peaks), SA2-only peaks did not appear to interact and form loops (**Fig. S4G**). We also asked whether SA2-only might make loops with any other STAG-bound sites; we used FitHiC (Ay et al. 2014) to call all loop-like interactions in our Hi-C matrices genome-wide, and then mined those with an SA2-only peak in at least one of its anchors. This only returned 126 loops of ∼0.8-Mbp in size that were also not sensitive to SA2 auxin-mediated degradation (**Fig. S4G**). In contrast, SA1-only-bound sites gave rise to >1600 long-range loops in *cis*, which were markedly weakened upon auxin-mediated SA1 degradation (**Fig. 4F**). Surprisingly though, ∼2000 loops emerge in SA1-depleted cells connecting previously-bound SA1-only sites (**Fig. 4F**). These were not seen as strong loops in non-auxin-treated cells, indicating widespread loop rewiring in the absence of SA1. Notably, these two subsets contain 833 loops with SA1-only anchors that persist upon SA1 degradation (**Fig. S5A**) and are on average shorter-range ones (**Fig. S5B**). These persistent SA1-only loops are significantly more enriched for convergent CTCF DNA-binding motifs at their anchors, compared to SA1-only loops detected in the presence or absence of SA1 (**Fig. S5C**). This is in line what was shown for strong loops across numerous mammalian cell types (Rao et al. 2014; Guo et al. 2015) co-bound by cohesin (Sofueva et al. 2013; Zuin et al. 2014a). Collectively, this analysis highlights reciprocal and subunit-specific contribution of SA1 and SA2 to 3D chromatin organization at different scales. At the Mbp-level SA1 depletion leads to more dramatic contact reshuffling than SA2, and this trend remains so at higher resolution, where individual looping events exclusively emanating from SA1-only sites are selectively lost or rewired upon SA1 depletion. On the other hand, SA2-only peaks do not have significant looping contribution, and might rather directly affect gene expression (as recently proposed; (Kojic et al. 2018).

### SA1 and SA2 depletion affects distinct gene subsets

Our analyses of genome-wide binding sites and their contribution to chromatin looping indicated that SA1 and SA2 might affect gene regulation via distinct paths. This prompted us to identify genes differentially-expressed upon depletion of either STAG subunit in HCT116 cells. We performed RNA-seq in two independent SA1-AID and SA2-AID clones. After filtering out genes responding to auxin treatment (Rao et al. 2017), we observed 167 and 169 strongly misregulated genes (FC>0,6; *P*-value <0.05) following SA1 and SA2 depletion, respectively (**Fig. 5A,B and S6A,B**). In each gene list we find only 36 genes that are reported to be misregulated upon RAD21 degradation in the same cell type (Rao et al. 2017) (**Fig. S6C**). Importantly, changes in gene expression levels for most of these genes are solely dependent on SA1 or SA2, and are not seen in the reciprocal AID cell line, as confirmed by RT-qPCR analysis for selected genes (**Fig. 5C**). This hints to specific roles for SA1 or SA2 in gene expression regulation. Gene ontology analysis with IPA (Ingenuity Pathway Analysis; (Kramer et al. 2014) revealed that SA1 and SA2 control genes belonging to different pathways and networks. For instance, SA1 depletion affected genes involved in “behavior”, “nervous system development”, and “cell-to-cell signaling and interaction” (**Fig. S6D**) in line with a proposed role for STAGs in neuronal development (Soardi et al. 2017). On the other hand, genes related to the development and function of cardiovascular and hematological tissues, as has been reported for SA2-deficient patients (Soardi et al. 2017) (Wessels et al., *manuscript in preparation*).

**Figure 5.**
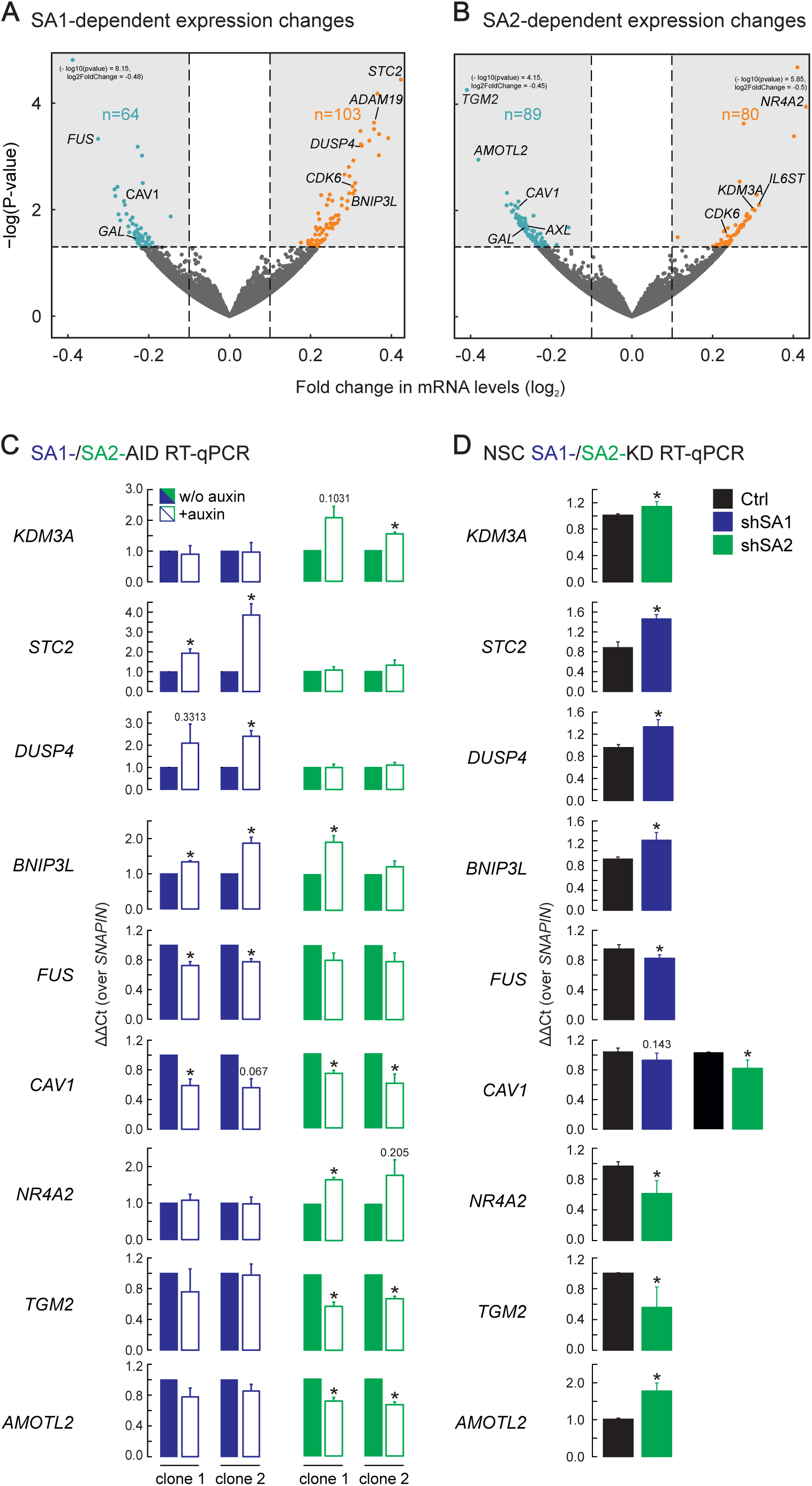
SA1 or SA2 regulate different genes. **(A, B)** Vulcano plots representing the transcriptional changes after SA1 degradation (A) or SA2 degradation (B). Significantly changing genes are plotted in green, genes that also respond to the degradation of the respective other STAG subunit in blue. Genes that lie outside of the plotted range are indicated with their fold change and p-value. **(C)** Validation of genes responding to degradation of SA1 or SA2 in two independent clones of the respective AID cells by RT-PCR/qPCR. Genes that respond to SA1 (STC2, DUSP4, BNIP3L, FUS) or SA2 (NR4A2, KDM3A, TGM2, AMOTL2) or both (CAV1) were tested. **(D)** Sensitivity of genes to depletion of SA1 or SA2 was recapitulated in neural stem cells using siRNA depletion of SA1 (ShSA1) and SA2 (ShSA2). (mean of n=3, T-test p-values are indicated, * P< 0,05; ** P< 0,005)

To assess the significance of the gene network predictions in a relevant cellular context, we used human neural stem cells (hNSCs) where SA1 or SA2 were sufficiently depleted by siRNA knockdown (**Fig. S7A, B**). Therein, we could confirm the dependency on SA1 or SA2 for a number of genes in this cell type, (**Fig. 5D**), while discrepancies could be due to differential gene dependencies between cell types. We also used RNA-seq data to select genes associated with microcephaly, like *CDK6* (Faheem et al. 2015), intellectual disability, like *AXL* (Burstyn-Cohen 2017), neural protection and development, like *IL6ST* (Marz et al. 1997), neurogenesis, like ADAM19 (Alfandari and Taneyhill 2018), or the neuro-regulatory peptide-encoding gene *GAL* (Borroto-Escuela et al. 2017) and tested whether they depend on cohesin in general or in particular on SA1 and SA2 for proper expression. We performed *SA1*, *SA2* or *SMC1A* knockdowns (**Fig. S7A-C**) and found that their expression in hNSCs is cohesin-dependent. Some genes, like *CDK6* and *ADAM19*, respond to the depletion of both SA1 and SA2, while others, like *AXL* and *IL6ST*, only to the loss of SA2 (**Fig. 6**). In summary, many of the gene expression changes seen in our HCT116 model translate into dependencies in a cell type relevant to neural development, with many of these genes also being documented for their involvement in cancer (Mochizuki and Okada 2007; Tigan et al. 2016).

**Figure 6.**
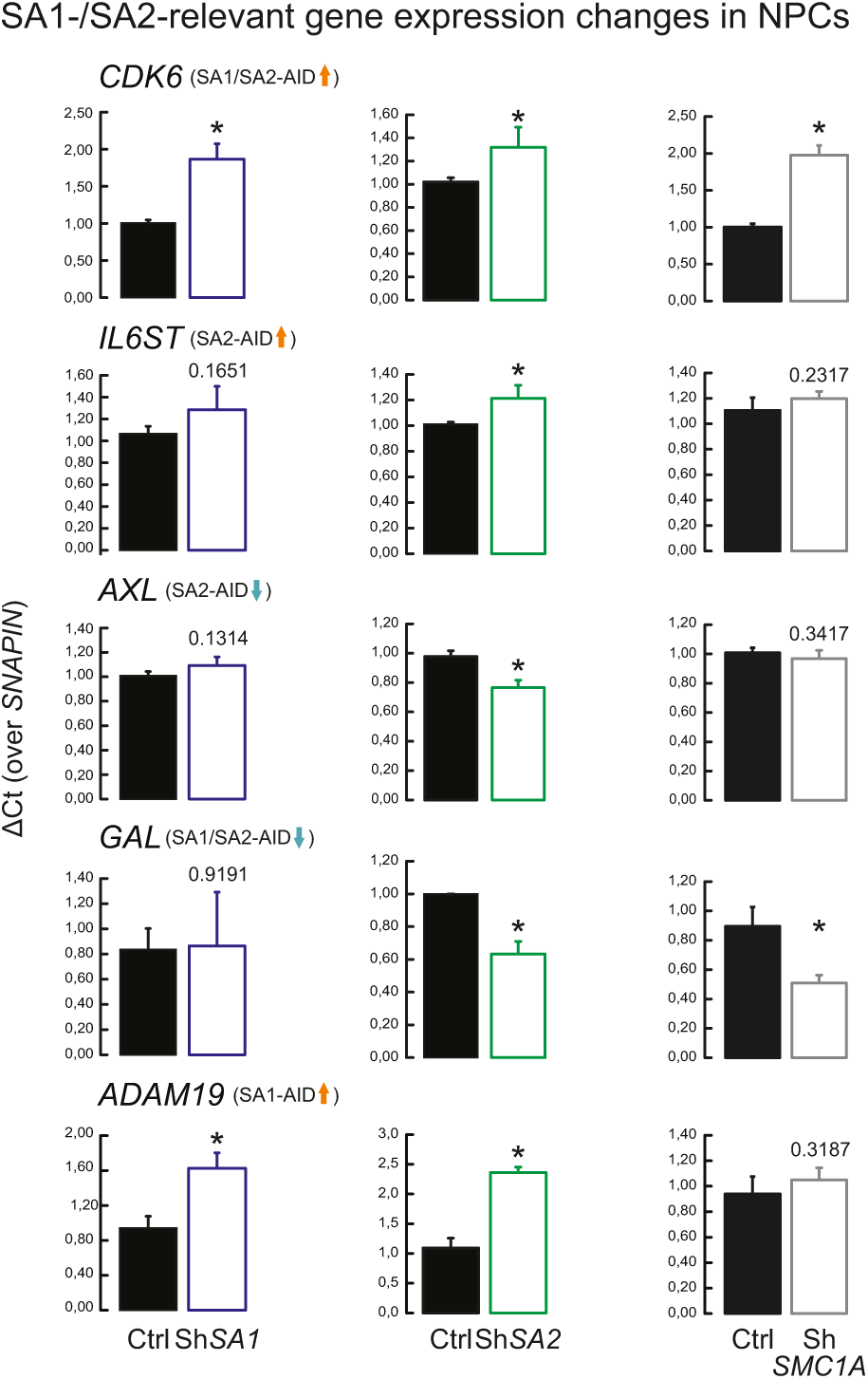
SA1 and SA2 regulate genes involved in neuronal development. Genes found to be involved in neuronal development and misregulated by SA1 and SA2 degradation were tested in neural stem cells under control (Ctrl), SA1 (ShSA1), SA2 (ShSA2) and SMC1A (ShSMC1A) siRNA knockdown to test whether they depend specifically on SA1 or SA2 or on the presence of the cohesin complex in general. (mean of n=3, T-test p-values are indicated, * P< 0,05; ** P< 0,005)

Finally, to understand how the depletion of SA1 or SA2 underpins misregulation of particular gene subsets, we plotted cohesin (RAD21, SMC1A) and CTCF ChIP-seq signal relative to the promoter (transcription start site, TSS) and the 3’ end of the gene (transcription termination site, TTS) and up to 6 kbp up- and downstream of SA1-/SA2-regulated genes (**Fig. S8A**). For genes downregulated after SA1 degradation, we observed enriched cohesin signals at the TSS, but not CTCF, pointing to a CTCF-independent binding here (as reported for particular promoters before; (Schmidt et al. 2010). Genes downregulated after SA2 degradation have enriched cohesin and CTCF signal at both their TSSs and TTSs, suggesting that long-range interactions spanning gene bodies might be involved in their regulation. Interestingly, genes downregulated upon SA2 depletion show a local accumulation of SA2 binding at their 3’ ends compared to the rest of the gene body and despite TSS-bound cohesin. On the other hand, all upregulated genes have cohesin and CTCF signals enriched at their promoters, and we speculated that these promoters are now contacting non-cognate enhancers due to a reshuffling of promoter-based long-range interactions, in line with the mechanism proposed to follow cohesin degradation (Rao, Huang et al. 2017). To this end, we plotted average Hi-C profiles around the TSS of these four SA1-/SA2-regulated gene subsets (**Fig. S8B**). For genes upregulated upon SA1 or SA2 depletion, we see that their TSS do not reside at a site of local insulation (i.e., at a TAD or loop domain boundary), but reshuffling of Hi-C contacts does occur at the longer- and shorter-range, respectively. This effect also holds true for downregulated genes, but here there is a stronger indication of them residing at a boundary site remodeled upon SA1 or SA2 depletion (**Fig. S8B**). This agrees with the early observation that such boundary sites are often marked by expressed gene promoters (Dixon et al. 2012) and their downregulation would then affect boundary strength.

## Discussion

Depletion of the cohesin STAG subunits is only lethal for cells if both SA1 and SA2 are knocked down (van der Lelij et al. 2017). The same is true for 3D chromatin organization: while RAD21 degradation eliminates all loops and TADs (Rao et al. 2017), degradation of SA1 or SA2 in our study does not lead to a complete breakdown of these structures. However, the analysis of our high resolution Hi-C data in respect to SA1-only, SA2-only and SA1/SA2-shared binding sites provides exciting insight into how SA1 and SA2 respectively contribute to chromatin folding.

While distinct roles for SA1 and SA2 have been proposed for their binding at telomeres and centromeres (Canudas and Smith 2009; Bisht et al. 2013), ChIP experiments for these two factors revealed extensive overlap between the two, as well as with CTCF along human chromosomal arms (Wendt et al. 2008; Remeseiro et al. 2012) (**Fig. 2A** and **S2**). Our ChIP-seq experiments exploited the mClover-tag in the *SA1* and *SA2* loci, thereby eliminating biases coming from the use of different antibodies. We again saw extensive overlap between SA1 and SA2 peaks, all of which align well to published RAD21 ChIP-seq data from HCT116 cells (Rao et al. 2017). However, we observed that 26% of all SA1 and 4% of all SA2 binding sites did not show binding by the other STAG subunit. These uniquely occupied sites are still occupied by CTCF, which contrasts previous observations of a large number of SA2-only peaks lacking CTCF (Kojic et al. 2018). This could be attributed to cell type-specific differences, since the relative numbers of SA1- and SA2-bound sites differ significantly between the cell types examined to date (Kojic et al. 2018). However, a very high >90 % overlap between SA2 and CTCF was already documented in HeLa cells, which contain substantially more SA2 than SA1 (Hauf et al. 2005; Wendt et al. 2008).

Interestingly, our Re-ChIP and single-molecule dSTORM localization experiments reveal that, at the level of the single alleles, individual binding sites are actually occupied exclusively by either cohesin-SA1 or cohesin-SA2 (again contradicting recent propositions of co-binding; (Kojic et al. 2018). The overlap of SA1 and SA2 ChIP-seq peaks then simply reflects the fact that these sites have no preference for SA1 or SA2 over the population average, and that SA1-binding might increase upon SA2 depletion and *vice versa* – an effect we clearly observe. SA1/SA2-shared sites, as well as SA1-only binding sites (but not SA2-only sites), display chromatin insulation properties (**Fig. 4G**). This is in agreement with the observation that SA1/SA2-shared sites have a similar genomic distribution as SA1-only sites, in contrast to SA2-only sites that preferentially localize to gene promoters.

Upon SA1- or SA2-degradation, functional differences between SA1 and SA2 become apparent also at their shared sites. SA2 depletion leads to a loss of insulation and to a mild, yet noticeable, increase in shorter-range interactions, visible in both the insulation (**Fig. 4G**) and the decay plots (**Fig. 4E**). In contrast, upon SA1 depletion we observe a reduction in interactions at the range of TAD sizes (i.e., up to ∼2 Mbp; **Figs. 4E and G**). This discrepancy can be interpreted on the basis of (i) the increase in SA1 occupancy at SA2-bound sites in the absence of SA2 that is more pronounced than the converse (**Fig. S1G-K**) and (ii) the combination of FRAP studies and loop extrusion modeling that suggest continuous formation and collapse of chromatin loops relying on cohesin and CTCF throughout the cell cycle (Hansen et al. 2017). In other words, introduction of more SA1 can lead to the establishment of new loops within Mbp-sized domains. This more dominant role for SA1 in loop formation is also supported by the reported lag between SA1 and SA2 association to chromatin after mitosis recorded in HeLa cells, which would suggest that loop extrusion by SA1 might commence earlier (Cai et al. 2018). Despite the robust and acute degradation of either STAG subunit in our cells, not all loops forming between SA1-only sites are sensitive to SA1 degradation. On one hand, there is a subset of ∼800 loops that persist despite SA1 depletion, and on the other >1,000 loops form *de novo* between previously SA1-only occupied sites. This likely indicates that replacing SA1 by SA2 at these sites suffices for loop formation, albeit not without reshuffling loop anchors since SA2-only sites form a strikingly small number of loops amongst them or with other SA1/SA2 sites (**Fig. S4F, G**). Thus, we suggest that context (i.e., the orientation of the underlying CTCF sites and the repertoire of co-bound factors at the anchors of prospective loops) is decisive for chromatin folding via SA1/2 and determines redundant and unique contributions.

Degradation of SA1 or SA2 also led to moderate changes in gene expression that concerned a limited number of genes (**Fig. 5**). However, expression of most of these genes was specifically affected only in response to the depletion of one or the other subunit, and was not observed upon RAD21 degradation (Rao et al. 2017) which should inactivate all cohesin complexes. This pattern was recapitulated in normal diploid cells using siRNA-mediated depletion of either STAG subunit, and is a strong indication of SA1-/SA2-specific dependencies. For SA2-dependent genes, given the preferential SA2 positioning at promoters and its general non-involvement in long-range looping, we suggest it confers direct gene regulation as has been suggested already (Kojic et al. 2018). For SA1-dependent genes, we saw that their TSSs generally resided at a position of local insulation based on Hi-C data, and thus deregulation of SA1-driven looping would reshuffle long-range contacts at this promoter and alter its regulation – an effect in line with what has been observed on RAD21 depletion (Rao et al. 2017). Notably, SA1-/SA2-specific gene expression dependencies we observed involve key developmental genes, like ADAM19, ZWINT, AXL and GAL.

Finally, our dSTORM imaging revealed – to our knowledge for the first time – that approximately only half of the observed CTCF clusters are in contact with cohesin in skin fibroblasts. Since in these cells the vast majority of CTCFs are chromatin-bound, CTCF clusters not contacting cohesin could represent the non-specific DNA-binding pool of CTCF observed by FRAP experiments in U2OS cells (Hansen et al. 2017); alternatively, it can be that only a subset of CTCF binding sites in a cell are occupied by cohesin at a given time. The size of the CTCF clusters then increases as the number of contacting cohesin cluster rises – for instance, the CTCF cluster sizes doubles when associated with two cohesin clusters. It is thus tempting to speculate that these larger clusters represent CTCF sites engaged in chromatin loops, although we did not directly visualize loops here. This hypothesis is reinforced by our observation that the median CTCF cluster size in RAD21-mAID HCT116 cells (Natsume et al. 2016a) decreases upon RAD21 degradation to a size comparable to that of the smallest CTCF clusters in skin fibroblasts. This rules out that larger CTCF clusters are merely neighboring sites and points to them being riggers that inform 3D chromatin structure, and which are most often found overlapping SA1-than SA2-cohesin clusters.

Taken together, we identified both redundant and non-redundant roles for the SA1 and SA2 subunits of the cohesin ring complex in looping and ultimately in gene regulation. Of course, much of the current knowledge on how these large protein complexes operate come from population studies complemented by (but not fully aligned to) single-cell measurements. Thus, in the future, it will be important to bridge this gap and address cell-to-cell variation in cohesin/CTCF function in genome organization.

## Materials and methods

### Cell lines

SA1-AID and SA2-AID cell lines were generated using the HCT116 CMV-OsTIR1 cells obtained from M. Kanemaki using the same strategy as described (Natsume et al. 2016a). In brief, to insert the mAID-mClover neomycin cassette (plasmid pMK289 obtained from Addgene) at the 3’ end of the SA1 and SA2 genes, 500-800nt homology arms were inserted left and right to the cassette present in the pMK289 plasmid. A gRNA targeting the 3’ end of the SA1 or SA2 gene in front of the stop codon was cloned into a Cas9 expression construct (Addgene p48193). Both, the gRNA/Cas9 plasmid as well as the plasmid carrying homology arms and mAID-mClover neomycin cassette were transfected into HCT116 CMV-OsTIR1 cells using FuGene HD (Promega). After 6 days selection with 700 µg/ml G418 the cells were FACS-sorted for mClover expression and seeded into 96 well plates to grow single clones. Screening of clones was performed based on nuclear localization of the mClover signal and PCR to identify homozygous clones. We noted that the expression of SA1-mAID-mClover (SA1-AID) and SA2-AID-mClover (SA2-AID) is somewhat lower than the wildtype protein. However, since the expression levels of SA1 and SA2 are quite variable between cell lines (Kojic et al. 2018) we concluded that this will not influence the planned experiments. To induce protein degradation 500 µM indole-3-acetic acid (auxin, IAA) dissolved in ethanol was added to the cells.

guide RNA:

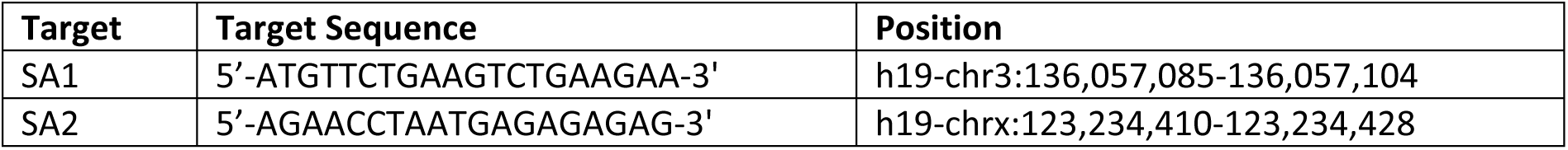

### Cell culture

SA1-AID and SA2-AID cells as well as RAD21-AID cells (Natsume et al. 2016b) were cultured in McCoy’s 5A medium supplemented with 10% FBS and PSG. For all auxin degradation experiments using SA1-AID and SA2-AID cells auxin or the solvent ethanol as control were added for 12 hours.

Human primary skin fibroblasts (Dept. of Clinical Genetics, Erasmus MC) were cultured in DMEM medium with 10% FBS and PSG.

### Antibodies

Rabbit polyclonal antibodies directed against EGFP, SA1 and SA2 were obtained by immunization of rabbits with either the full-length protein (EGFP) or N-terminal human protein fragments (SA1 – residues 1-73, SA2-residues 1-68) that were expressed as fusion protein with 6XHis-tag in E. coli and purified with NiNTA matrix (Qiagen). Immunization of the rabbits was performed by Absea (Beijing, China). Antibodies were purified from the serum using the proteins used for immunization coupled to sulfolink (Thermo Fisher) matrix. The anti-SMC3 antibody has been described before (Zuin et al. 2014b). Additional antibodies are listed in Table S1.

### Immunoprecipitation

Immunoprecipitation experiments were performed as described (Watrin et al. 2006)

### Cell fractionation

Whole-cell extracts of human primary skin fibroblasts were separated into soluble supernatant and chromatin-containing pellet fractions as described (Watrin et al. 2006).

### ChIP sample preparation

ChIP was performed as described (Cabianca et al. 2012) with some adaptations. Approximately 5*10^7^-1*10^8^ cells were crosslinked by replacing the medium with the same volume 1% formaldehyde in PBS (methanol-free, 16% Thermo Scientific) for 10 minutes at room temperature. After quenching with 0.125M Glycine (final concentration), cells were washed three times with PBS, harvested using a silicon scraper. Lysis of the cells was carried out by incubating the cells in LB1 buffer (50 mM Hepes-KOH, pH 7,5, 140 mM NaCl, 1 mM EDTA, 10% glycerol, 0,5% NP-40, 0,25% Triton X-100) for 5 minutes, followed by centrifugation at 1,350g for 5 min and incubation with LB2 buffer (10 mM Tris-HCl, pH= 8.,, 200 mM NaCl, 1 mM EDTA, 0,5 mM EGTA) for 10 minutes. Nuclei were pelleted down by centrifugation of the cells at 1,350g for 5 min and were resuspended in LB3 buffer (10 mM Tris-HCl, ph=8,0, 100 mM NaCl, 1 mM EDTA, 0,5 mM EGTA, 0,1% Na-Deoxycholate, 0,5% N-lauroylsarcosine). All lysis buffer contained 1x protease inhibitors. This protocol ensures the removal of cytoplasmic proteins from the lysates. Sonication was performed using a Diagenode bioruptor (15 min, 5-7 cycles, 30” each). Sonicated lysates were cleared by the addition of Triton X-100 to a final concentration of 1% and by centrifugation at maximum speed for 10 min at 4°C. 50 µl of the sonicated lysates were reserved for input and the remaining material stored at – 80°C until use. To prepare the antibody-loaded beads, 100 µl of Dynabeads were washed three times with blocking solution (1x PBS, 0,5% BSA) and incubated overnight with 10 µg of the required antibodies. For each ChIP 100 µg chromatin was incubated overnight at 4°C with the antibody-bound beads. Beads were washed 6X with RIPA buffer (50 mM Hepes-KOH, pH= 7,6, 500 mM LiCl, 1 mM EDTA, 1% NP-40, 0,7% Na-Deoxycholate), 1X with TE/50mM NaCl. The DNA was eluted from the beads by incubation with 250 µl of elution buffer (1x TE, 2% SDS) for 15 min at 65°C. Crosslinking was reverse by incubation at 65°C overnight, proteins were digested by incubation with 8 µl of 10 mg/ml of Proteinase K at 55°C for 1 hour and RNA degraded by incubation with 8µl of 10 mg/ml at 37°C for 30 min. DNA was purified then purified using phenol-chloroform extraction followed by ethanol precipitation. The DNA was then resuspended in 25 µL TE and used for either qPCR or sequencing. For indicated experiments the One Day ChIP kit (Diagenode) was used according to the protocol of the manufacturer.

The samples were either analyzed by qPCR, the respective primers are listed in Table S3, or processed for sequencing.

### ChIP-seq sample preparation and sequencing

NGS libraries have been prepped from the ChIP DNA using the NEXTFlex ChIP-Seq kit from BioO Scientific and sequenced according to the Illumina TruSeq Rapid v2 protocol on the HiSeq2500 for a single read 50bp in length and a 8bp dual index read. The data were de-multiplexed and mapped against GRCh38 using initially HiSat2 (Kim et al. 2015).

### ChIP-seq analysis

ChIP-seq data was aligned to hg19 using Bowtie (Langmead et al. 2009) under default settings and peak calling was carried out using MACS2 (Liu 2014)(Liu, 2014) setting a q value to 0.06 or 0.005. All data was normalized against the corresponding input control using the ‘-c’ option of MACS 2.0. Signal tracks were generated with the “bamCoverage” function of deepTools 3.1.3 (Ramirez et al. 2014) and were normalized to sequencing depth.

Cohesin positions (common, SA1-only and SA2-only sites) were defined by calculating the overlap between SA1 and SA2 peaks using the “intersect” option of BEDtools v2.27 (Quinlan and Hall 2010) with a minimum of 1 nt overlap. STAG1 peaks called using a p-value of 0.005 were intersected with STAG2 peaks called using a p-value of 0.06 to define STAG1 only sites. For STAG2 only sites, a list of STAG2 peaks (p-value = 0.05) was intersected with a list of STAG1 sites (p-value = 0.06).

Mean read density profiles and read density heatmaps were obtained with deepTools using normalized bigwig files and plotting them around peak summits of SA1 or SA2 only or common peaks. Enrichment of common and only sites at enhancer and promoters was calculated using the intersect function from BEDtools with a minimum overlap between peaks of 50%. Published ChIP-seq data for different histone modifications in HCT116 cells was used to define the enhancer and promoter regions. Enhancers were defined by the presence of H3K4me1, while active promoters were defined by the presence of H3K4me3 and the proximity to TSS (+/- 2Kb). A particular genomic region could only belong to one category. The presence of H3K27ac at these regulatory regions was also analyzed to identify active enhancers and promoters.

### Transcription factor mapping

The analysis of transcription factor motifs at SA1 and SA2 binding sites in context with DNase hypersensitivity as marker for the accessibility of these predicted TF sites was performed as described previously (Lin et al. 2015).

### Culturing of neural stem cells preparation and siRNA knockdown

The human neural stem cells (NSCs) were obtained from ThermoFisher/Invitrogen (N7800-100) and cultured in KnockOut™ D-MEM/F-12 medium (ThermoFisher 12660012) supplemented with 2mM L-glutamine (Gibco 25030081), 20 ng/ml EGF (Peprotech 315-09), 20 ng/ml bEGF (Peprotech 100-18B) and StemPro® Neural Supplement 2% (ThermoFisher A1050801).

The siRNA knockdown was performed by electroporation using the Amaxa nucleofector I in combination with the Amaxa Cell Line Nucleofector Kit V (Lonza, VCA-1003). SiRNA constructs with the siRNA cloned in the pLKO.1-Puro vector were obtained from the MISSION shRNA library (Sigma product SHGLY). The specific siRNA sequences used are:

Control  5’-CAACAAGATGAAGAGCACCAA-3

SA1   5’-CGTCGCTTTGCCCTTACATTT-3’

SA2   5’-GCAAGCAGTCTTCAGGTTAAA-3’

SMC1A  5’-CCAACATTGATGAGATCTATA-3’

In brief, about 3,5 million hNSCs were resuspended in 100µl supplemented transfection buffer with 3,5µg of DNA and electroporated with protocol A-33. The cells are then transferred with warm hNSC medium to a 6-well plate coated with Geltrex (Gibco, A1413202, incubate for 1hr at 37C). The day after transfection the medium was refreshed and 1µg/ml puromycin (Sigma, P8833) added. Cells were harvested 48 hours (SMC1A) and 72 hours (STAG2) after transfection.

### Transcription analysis by reverse transcription (RT) and qPCR

Cells were harvested and total RNA was prepared using Trizol Reagent (Invitrogen). RNA was purified with RNeasy Mini Kit according to manufacturer’s instructions and eluted in DEPC water. cDNA was generated by reverse transcription using oligo(dT)18 primer (Invitrogen), Superscript IV Reverse Transcriptase (RT) (Invitrogen) and RNaseOUT Recombinant Ribonuclease Inhibitor (Invitrogen) according to the manufacturer’s instructions. The amounts of the different transcripts were compared by qPCR using SYBR Green and Platinum Taq Polymerase (Invitrogen) in CFX96 light cycler (BioRad) and specific primers. The ΔΔCt method was used to calculate the fold change in gene expression using the housekeeping gene *SNAPIN* and the control sample for normalization. The sequences of the primers used are listed in Table S3.

### Re-ChIP or sequential ChIP

The original protocol from Voelkel et al. (Volkel et al. 2015)was modified to perform the 1^st^ ChIP step with antibodies crosslinked to beads. First, 20 mg of antibody were loaded on 50 µl Affi-Prep® Protein A Resin (Biorad) (multiply this with the number of 2^nd^ CHIP’s planned and plan one sample to check the ChIP efficiency of the 1^st^ ChIP step). After three washes with TBS/T (0,01%Triton X100) the beads were washed 3 times with 0,2M Sodiumborate pH 9,0 and then incubated for 20 min with a 20 mM solution of Dimethyl pimelimidate dihydrochloride (Sigma Aldrich) at room temperature under rotation. The crosslinking reaction was quenched by washing the beads three times with 250 mM Tris pH 8. Not-crosslinked antibodies were removed by a short treatment with 50 µl 100mM Glycine pH 2.0 per 50 µl beads and then the beads are neutralized again by washing with TBS/T. The 1^st^ ChIP was performed as described in the protocol for ChIP-sequencing with the exception that the beads were eluted twice with (25 µl per 50 µl beads)100 mM NaHCO3, 1% SDS, 10 mM DTT for 30 min at 37 °C. The eluates were diluted 1:50 with ChIP buffer from the One Day ChIP kit (Diagenode) and subsequently subjected to a second ChIP in accordance with the One Day ChIP kit manual but performing an overnight antibody incubation under rotation at 4°C. Primer sequences for ChIP-qPCRs are listed in Table S2.

### RNA sample preparation and sequencing

RNA-Seq libraries were prepared according to the Illumina TruSeq stranded mRNA protocol (www.illumina.com) starting from 200ng total RNA. One microliter of library was loaded on an Agilent Technologies 2100 Bioanalyzer using a DNA 1000 assay to determine the library concentration and for quality check. The libraryes were sequenced according to the Illumina TruSeq Rapid v2 protocol on the HiSeq2500 for a single read 50bp in length and a 8bp dual index read. The data were de-multiplexed and Illumina adapter sequences have been trimmed off the reads, which were subsequently mapped against the GRCh38 human reference using HiSat2 (version 2.1.0) (Kim et al. 2015) initially.

### RNA-seq analysis

Raw reads were mapped to the human reference genome (hg19) using default settings of the STAR aligner (Dobin et al. 2013), followed by quantification of unique counts using *featureCounts* (Liao et al. 2014). Counts were further normalized via the RUVs function of RUVseq (Risso et al. 2014) to estimate factors of unwanted variation using those genes in the replicates for which the covariates of interest remain constant and correct for unwanted variation, before differential gene expression was estimated using DESeq2 (Love et al. 2014). Genes with an FDR <0.05 and an absolute (log_2_) fold-change of >0.6 were deemed as differentially-expressed and listed in Tables S4 and S5. Volcano plots were generated using R (http://www.R-project.org). Significant deregulated genes (p-value > 0,05) were labeled in green. Genes found to be deregulated in both cases, after depletion of SA1 and SA2, where labeled in blue. Gene ontology and networks were generated through Metascape (Zhou et al. 2019).

### Hi-C sample preparation, sequencing, and data analysis

*In situ* Hi-C data from SA1-/SA2-AID HTC116 cells were generated as described previously (Zirkel et al. 2018) in two biological replicates each. Resulting DNA libraries were paired-end sequenced to ∼300 million read pairs each on a HiSeq4000 platform (Illumina). Raw reads were mapped, annotated, and corrected for biases using the Juicer suite (Durand et al. 2016b), before interactive visualization via Juicebox (Durand et al. 2016a). After ensuring replicate reproducibility using HiCRep (Yang et al. 2017), data from the same treatments were merged for all downstream analyses. Compartment analysis was performed using the first principal component of the Hi-C matrices as described (Rao et al. 2017) and TAD boundaries were identified using default setting in (Kruse et al. 2016). Binarization and matrix subtractions were performed as previously described (Zirkel et al. 2018) and custom scripts are available on request. For plotting insulation heatmaps and averaged loop profiles, normalized interactions values in the twenty 10-kbp bins around each SA1 or SA2 peak were added up, normalized to the median value in each matrix and plotted provided the local maxima are higher than the third quantile of all Hi-C data in the matrix. For the decay plots, matrices generated in JUICER (Durand et al. 2016b) that were normalized with the Knight-Ruiz matrix balancing normalization. All *in house* programming scripts used for this analysis are available on request.

### Immunostaining (including dSTORM preparation)

Cells were seeded to 70% confluency at the time of fixation on cover slips. Fixation was performed with 4% paraformaldehyde in PBS pH 7.4 for 20 min at room temperature. After fixation, cells were washed three times with ice-cold PBS and were permeabilized with 0.1% Triton X-100 in PBS for 5 minutes (8 min for fibroblasts) at room temperature. Cells were blocked with 3% BSA in PBS/T for 1 hour. SA1-AID and SA2-AID cells were incubated for 1 hour with a-EGFP and after washing goat anti rabbit Alexa 488. Coverslips were mounted with Prolong-Gold and imaged with a Zeiss Axio Imager Z1 Apotome microscope (Carl Zeiss, Germany). For dSTORM cover slips were incubated with the primary antibodies (goat-antin-SA1, Abcam; mouse-anti-CTCF, BD); and rabbit-antiSA2, Bethyl labs) over night at 4 degree and after washing with the secondary antibodies (donkey-anti-rabbit Alexa647, donkey-anti-goat Alexa488 and donkey-anti-mouse CF568) for 45 min at room temperature.

### dSTORM imaging

Cell were seeded to 70% density on cover slips, fixed and stained cells were mounted in an attofluor^TM^ cell chamber (Thermo Fisher) and and 1ml of dSTORM buffer (25mM MEA (Sigma), Glucose Oxidase (Sigma), Catalase (Sigma) 50mM NaCl and 10% Glucose (Sigma) in 10mM Tris-HCl pH 8). The cell chamber was sealed with a coverslip and incubated on the microscope at room temperature for 30 min prior to imaging, to minimize drift. Imaging was performed using a Zeiss Elyra PS1 system fitted with an Andor iXon DU 897, 512×512 EMCCD camera. Images were made using a 100x 1.49NA TIRF objective and were imaged in HiLO mode. High Power 100 mW diode lasers with wavelengths of 488, 561 and 642nm were used to excite the fluorophores and respectively BP495-575+LP750, BP 570-650+LP750 or LP655 filters were used. Movies of 12000 frames were recorded with an exposure time of 33ms. Multi-channel images were acquired sequentially from high wavelength to lower wavelengths.

### dSTORM analysis

Three dSTORM movies were made one for each protein and analyzed using Zeiss ZEN 2012 software. Localizations with a precision of larger than 50 nm were discarded, remaining localizations were drift corrected using a model-based approach. All additional analysis was done in R (http://www.R-project.org) using the SMoLR package (Paul et al. 2019) and FIJI (Schindelin et al. 2012). Localizations from a single nucleus were selected manually by selecting a region of interest (ROI) in FIJI and the IJROI_subset function in SMoLR. Localizations were clustered based on their density using a Kernel Density Estimation (KDE) based clustering algorithm with the threshold set to 0.05 for all three channels. All clusters for CTCF were selected and their area was measured using the thresholded KDE binary image. In the triple channel dSTORM experiments CTCF was grouped based on the number of overlapping binary clusters from the STAG1 and STAG2 channels, clusters were considered overlapping if pixels containing both colors are present in the structure.

## Acknowledgements

Work in the lab of K.S.W. was funded by the Dutch Cancer Society (KWF) grant EMCR 2015-7857 and by the by the Netherlands Organisation of Scientific Research (NWO-BBOL) grant 737.016.014. Work in the lab of A.P. was funded by the German Ministry for Research (DFG) via the PA2456/5-1 and SPP2202 grants and by CMMC JRGVIII core funding.

We thank Masato T. Kanemaki, National Institute of Genetics, Tokyo (Japan), for the HCT-116 CMV-OsTIR1 and HCT-116 RAD21-mAiD-mClover cells. We thank Jan-Michael Peters, IMP Vienna (Austria), for the rabbit-anti SA1 and rabbit-SA2 antibodies and Niels Galjart, Dept. of Cell Biology of the Erasmus MC, for the rat-anti SMC1A antibodies. We thank Karin Wisse for initiating the dSTORM experiments and re-ChIP experiments. We thank Mike R. Dekker and Raymond A. Poot for help with the siRNA knockdown in hNSC’s. We thank Yvonne Mueller, Department of Immunology of the Erasmus MC, for her help with FACS sorting. We would also like to thank Yulia Kargapolova and Matthew Ploetzke for help and advice on data analysis.

## AUTHOR CONTRIBUTIONS

VC JAS AP KSW conceived and designed the experiments. VC established the AID cell lines. VC, MMG, JAS, AZ and EO performed experiments. VC, MMG, EGG, JAS, NJ, AP analyzed the data. WvIJ and ABH contributed reagents, materials or analysis tools. AP and KSW jointly supervised research. VC, AP and KSW wrote the manuscript.

## Supplementary figures

**Figure S1.**
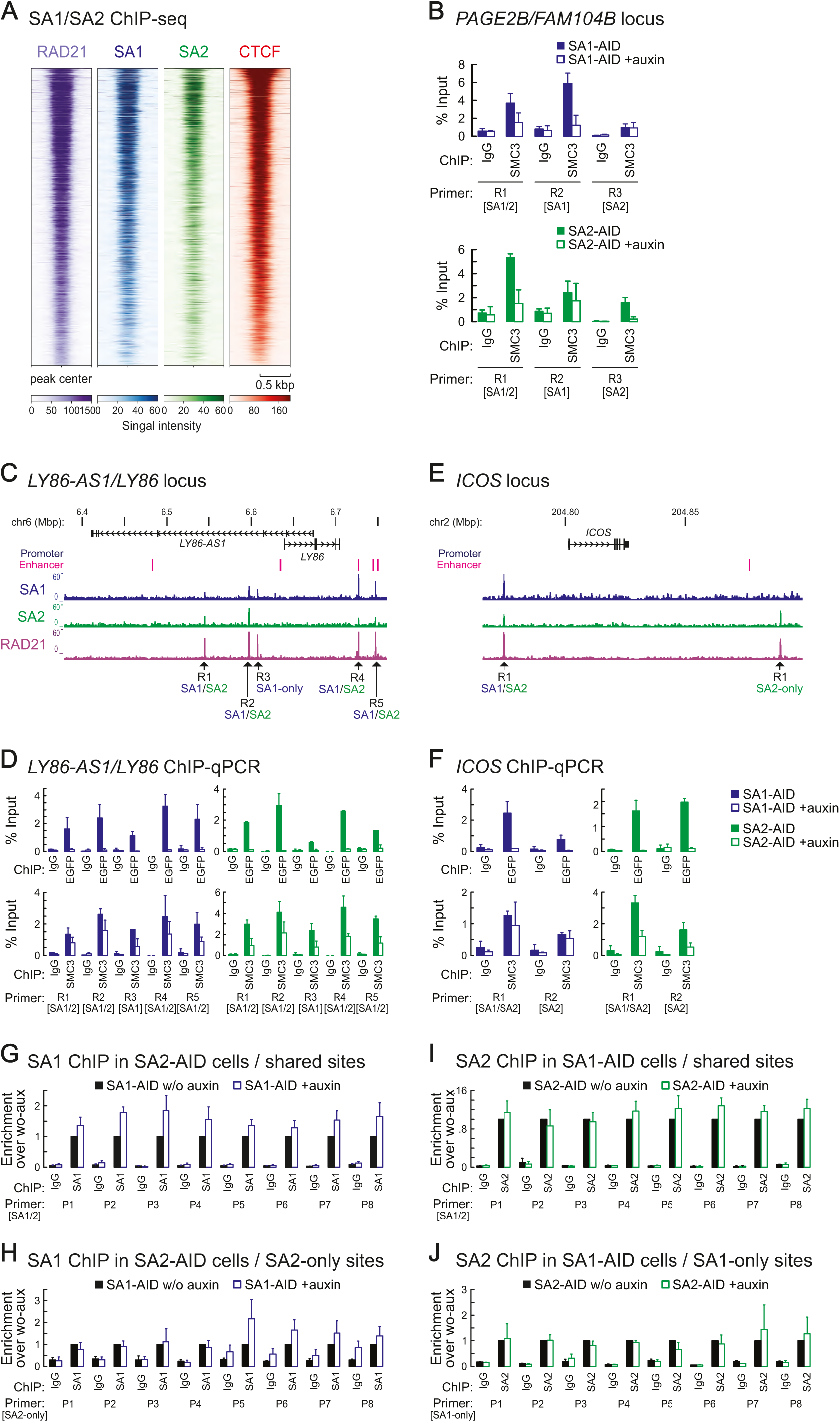
SA1 and SA2 have overlapping as well as discrete binding sites. **A)** Heatmaps showing the ChIP-seq signals for SA1, SA2, Rad21 and CTCF. RAD21 and CTCF ChIP-seq data are from Rao et al., 2017 (Rao et al. 2017). **B**) ChIP-qPCR showing that SMC3-binding at sites in the PAGE2B/FAM104B locus (also in Figure 2D, E) is reduced upon degradation of SA1 or SA2 on SA1- or SA2-only sites but is also on SA1/SA2-shared sites. **C, E**) Two more loci were analyzed to confirm the presence of SA1 and SA2-only sites, the LY86-AS1/LY86 locus (C) and the ICOS locus (E). **D, F**) SA1-only, SA2-only and SA1/2 shared sites at the LY86-AS1/LY86 locus (D) and the ICOS locus (F) were tested in anti-EGFP ChIP (upper panel) and SMC3 ChIP from SA1-AID and SA2-AID cells w/o and with auxin treatment. **G, H**) SA1 ChIP was performed from SA2-AID cells with and without auxin treatment to test whether SA1 accumulates more at these sites after SA2 depletion. SA1/SA2 shared sites (G) and SA1-only sites (H) were analyzed. **I, J**) SA2 ChIP was performed from SA1-AID cells with and without auxin treatment to test whether more SA2 accumulates more at these sites after SA1 depletion. SA1/SA2 shared sites (I) and SA2-only (J) sites were analyzed.

**Figure S2.**
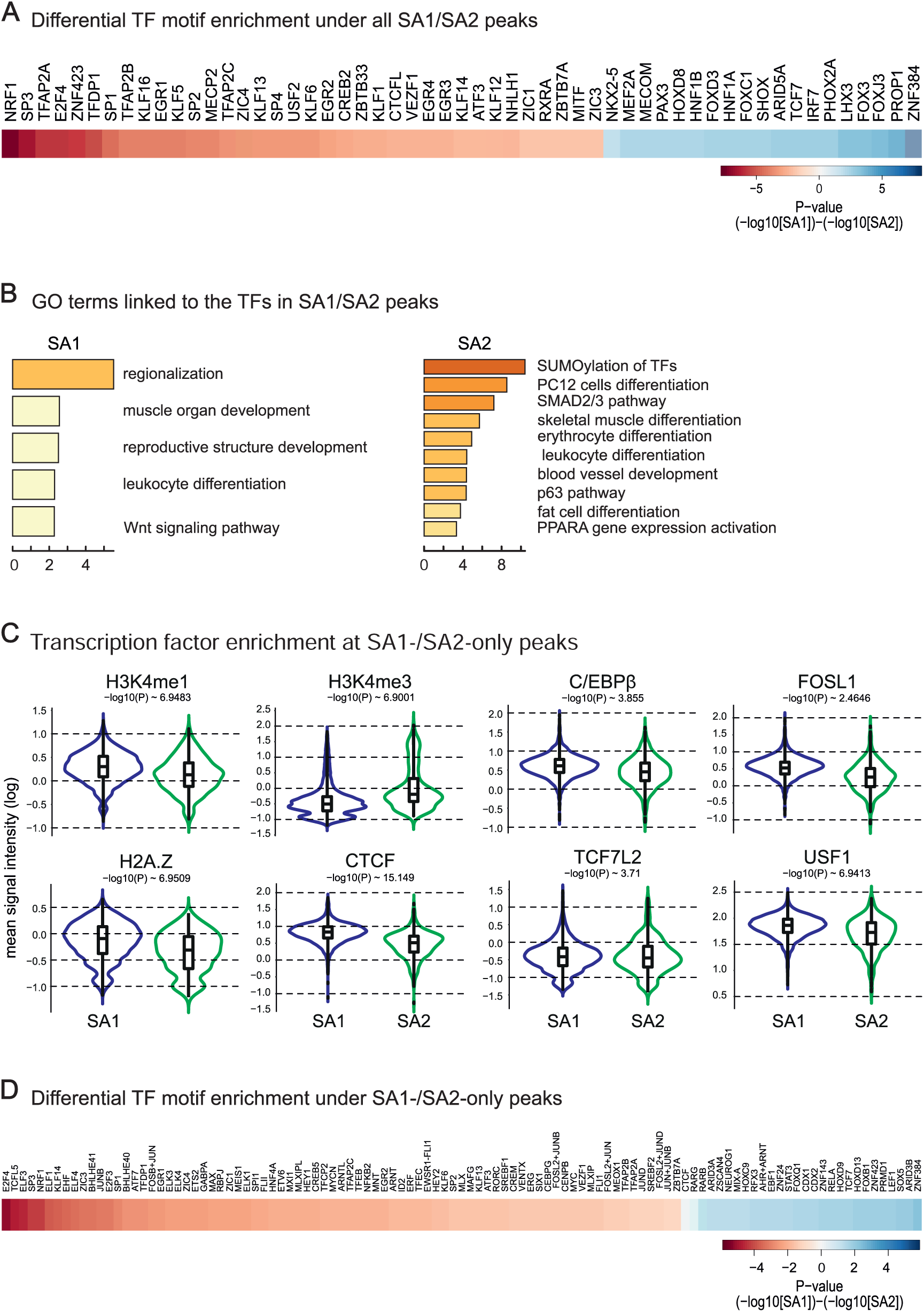
Enrichment of transcription factor motifs and signals at SA1- and SA2-only sites. **A)** The probability of the presence of another transcription at the SA1/SA2-shared binding sites was calculated based on the accessibility of the sites (DNase hypersensitivity) and the presence of transcription factor motifs (Lin et al. 2015). The differences of enrichment between SA1 and SA2 are plotted as heatmap. **B)** GO-term analysis of the transcription factors plotted in (A) using Metascape (Zhou et al. 2019). **C)** Enrichment of histone marks and transcription factors at SA1- and SA2-only peaks. Published datasets for HCT116 cells were used (Table S8). **D)** The probability of the presence of another transcription at the SA1- and SA2-only binding sites was calculated based on the accessibility of the sites (DNase hypersensitivity) and the presence of transcription factor motifs (Lin et al. 2015). The differences of enrichment between SA1 and SA2 are plotted as heatmap.

**Figure S3.**
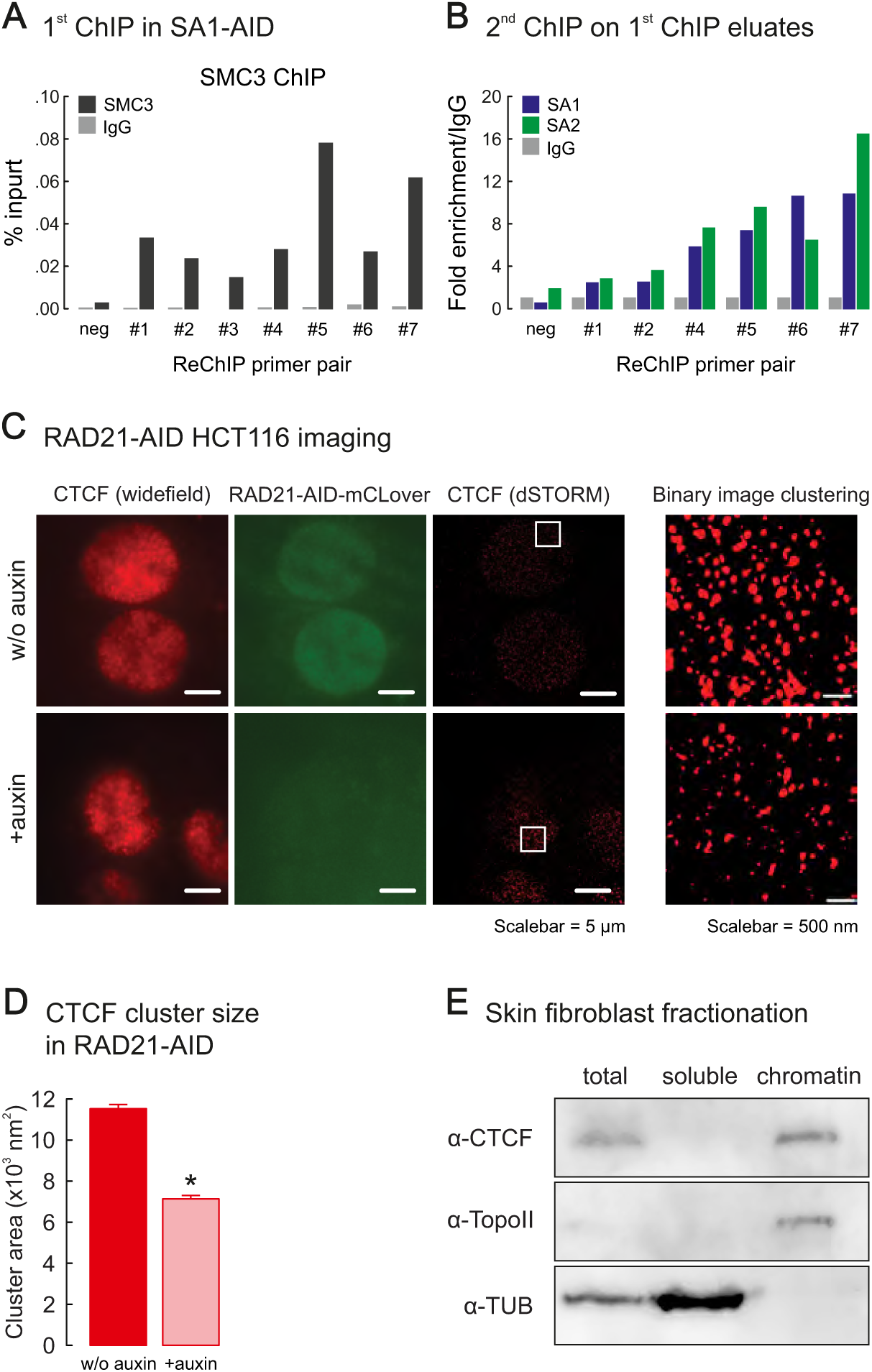
SA1 and SA2 do not co-occupy individual binding sites. **A**) As positive control for the Re-ChIP experiment, ChIP was performed using SA1AID cells with SMC3 antibodies crosslinked to the beads and analyzed with qPCR primers directed against one negative and seven positive cohesin binding sites. **B**) The eluate from the SMC3 ChIP was used for a second ChIP with anti-SA1 and anti-SA2 antibodies and analyzed with the same primers as in (A). **C**) dSTORM using anti-CTCF was performed in HCT116 RAD21-AID cells (Natsume et al. 2016a) w/o and with auxin treatment for 6 hours and the size of the CTCF signal clusters analyzed as in Figure 3D. **D**) The size of the CTCF clusters decreases after RAD21 degradation (+/- standard deviation of mean - SEM). **E**) Fractionation of human skin fibroblasts used for dSTORM in Figure 3 in soluble fraction and chromatin-bound fraction. Western blotting with anti-CTCF shows that CTCF is not detectable in the soluble fraction and therefore fully chromatin-bound. As controls for the chromatin-bound fraction a-Topoisomerase II was used and a-tubulin as control for the soluble fraction.

**Figure S4.**
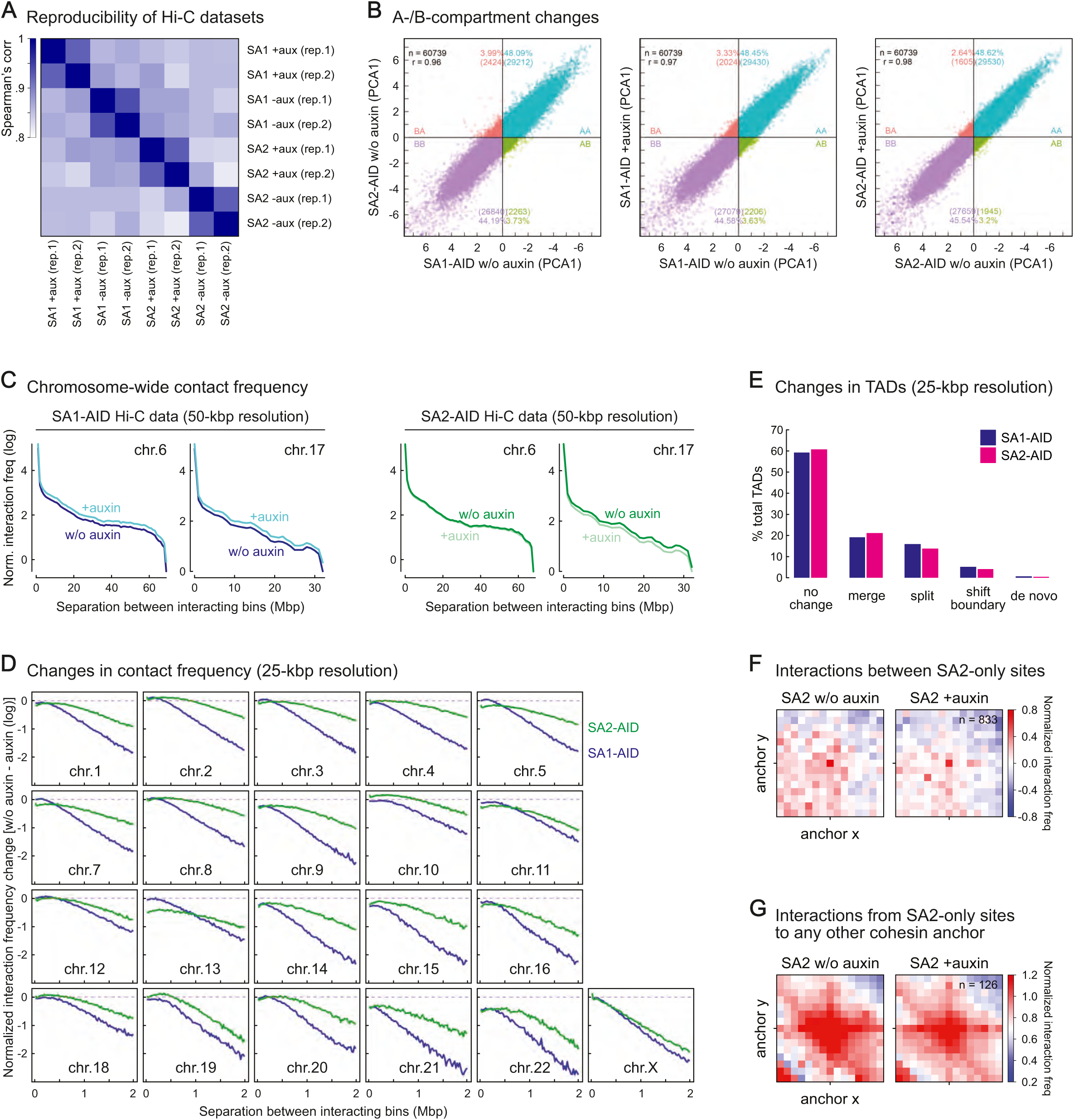
SA1 and SA2 affects long-range interactions differently. **A)** Correlation between the different samples and replicates of the Hi-C experiments in this study. **B)** Contacts between and within A and B compartments are correlated between the contact matrices of SA1-AID and SA2-AID cells without (w/o) auxin and then with and without auxin SA1-AID and SA2-AID cells. Total contacts observed, correlation coefficient as well as the percentage and number of the contacts between the specific compartments are indicated. **C)** Normalized interaction frequencies are plotted against the distance between interacting bins (50kb) without and with auxin-mediated degradation for SA1-AID and SA2-AID cells. Chromosomes 6 and 17 are shown. **D)** Change of contacts between bins (25kb) relative to the separation of the contacting fragments after SA1 or SA2 degradation are shown for all chromosomes, except chromosomes 6 and 17 that are displayed in Fig. 4E. **E)** Comparison of TADs before and after degradation of SA1 or SA2. **F**) Aggregate peak analysis of SA2-only peaks showing that SA2-only peaks do not contact each other. **G)** Aggregate peak analysis of the contacts of SA2-only peaks to other cohesin sites showing that these contacts persist after SA2 degradation.

**Figure S5.**
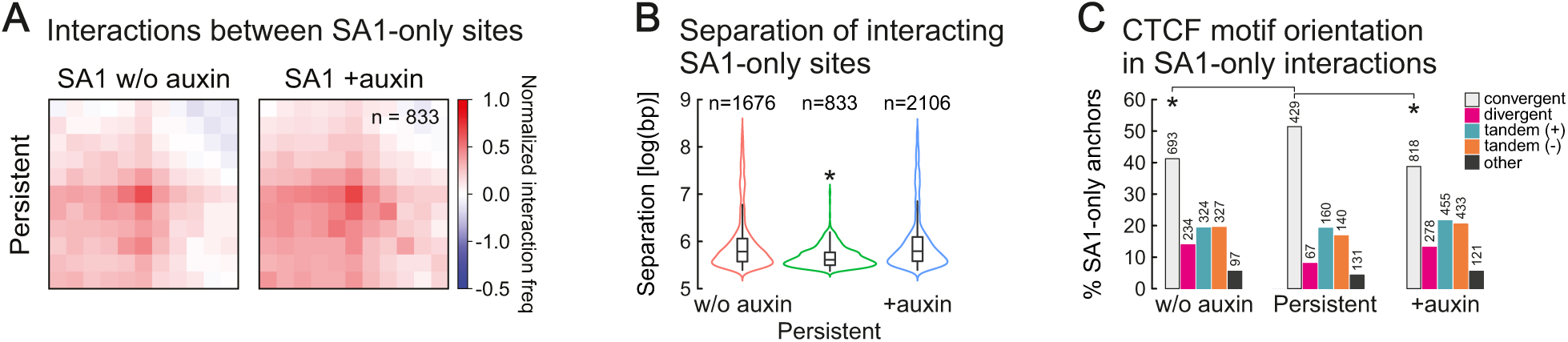
Contacts between SA1-only sites. **A)** Aggregate peak analysis of persistent contacts between SA1-only sites without (w/o) and with auxin in SA1-AID cells. **B)** Number and distances between interacting bins with SA1-only sites with contacts in SA1-AID cells without (w/o) auxin, with auxin (+auxin) and persistent in both situations. **C)** Orientation of the CTCF motif at SA1-only sites engaged in the different contact categories shown in (B) without (w/o) auxin, with auxin and persistent.

**Figure S6.**
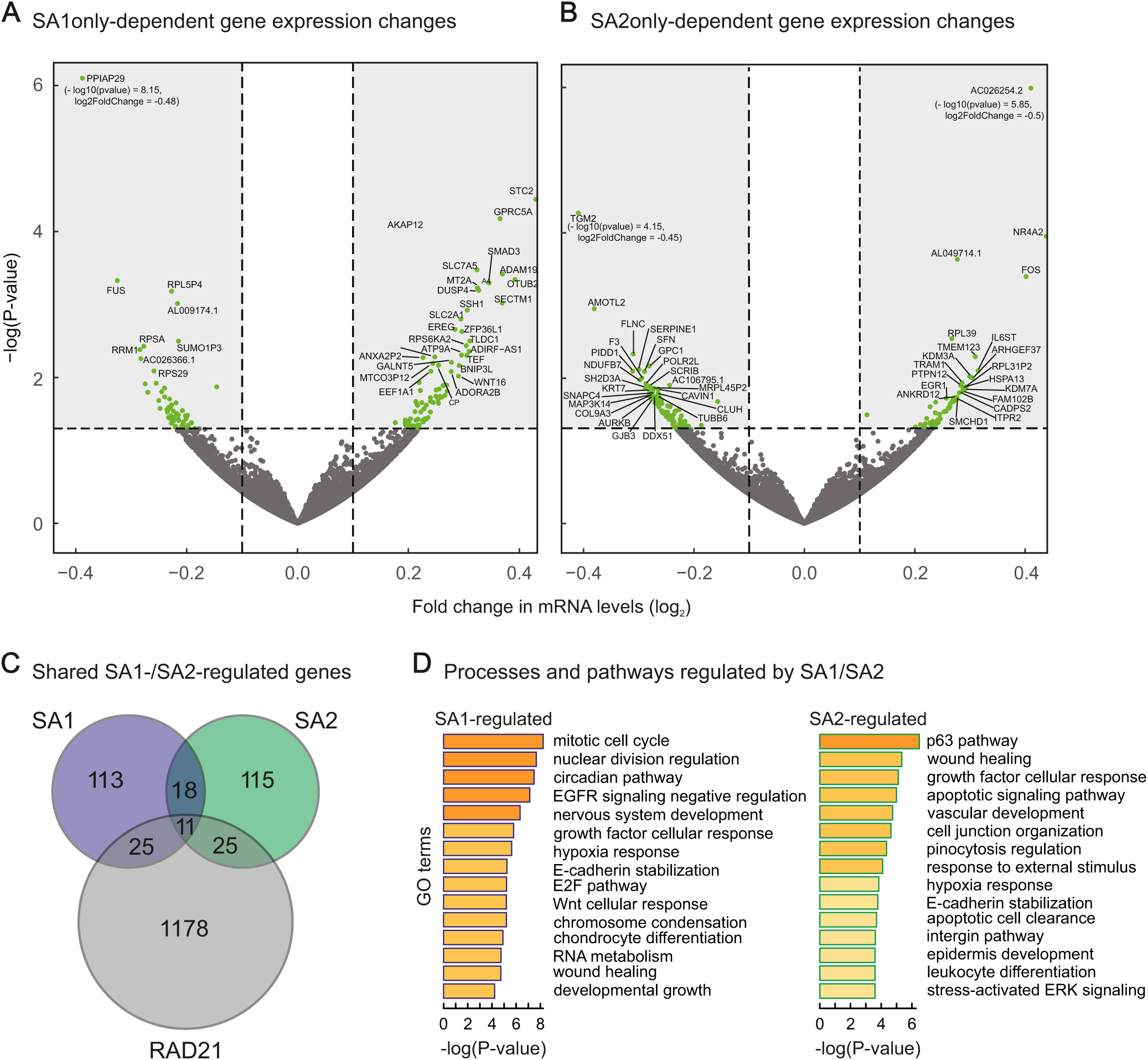
SA1 or SA2 regulate different genes. **A+B**) Vulcano plots representing the transcriptional changes after SA1 degradation (A) or SA2 degradation (B), presented without genes overlapping between the datasets. Significantly changing genes are plotted in green. Genes that lie outside of the plotted range are indicated with their fold change and p-value. **C**) Pie chart showing the overlap of the genes differentially expressed after SA1, SA2 and RAD21 degradation (Rao et al. 2017). **D**) Functional analysis of the genes differentially expressed after SA1 (STAG1) or SA2 (STAG2) degradation using Metascape (Zhou et al. 2019).

**Figure S7.**
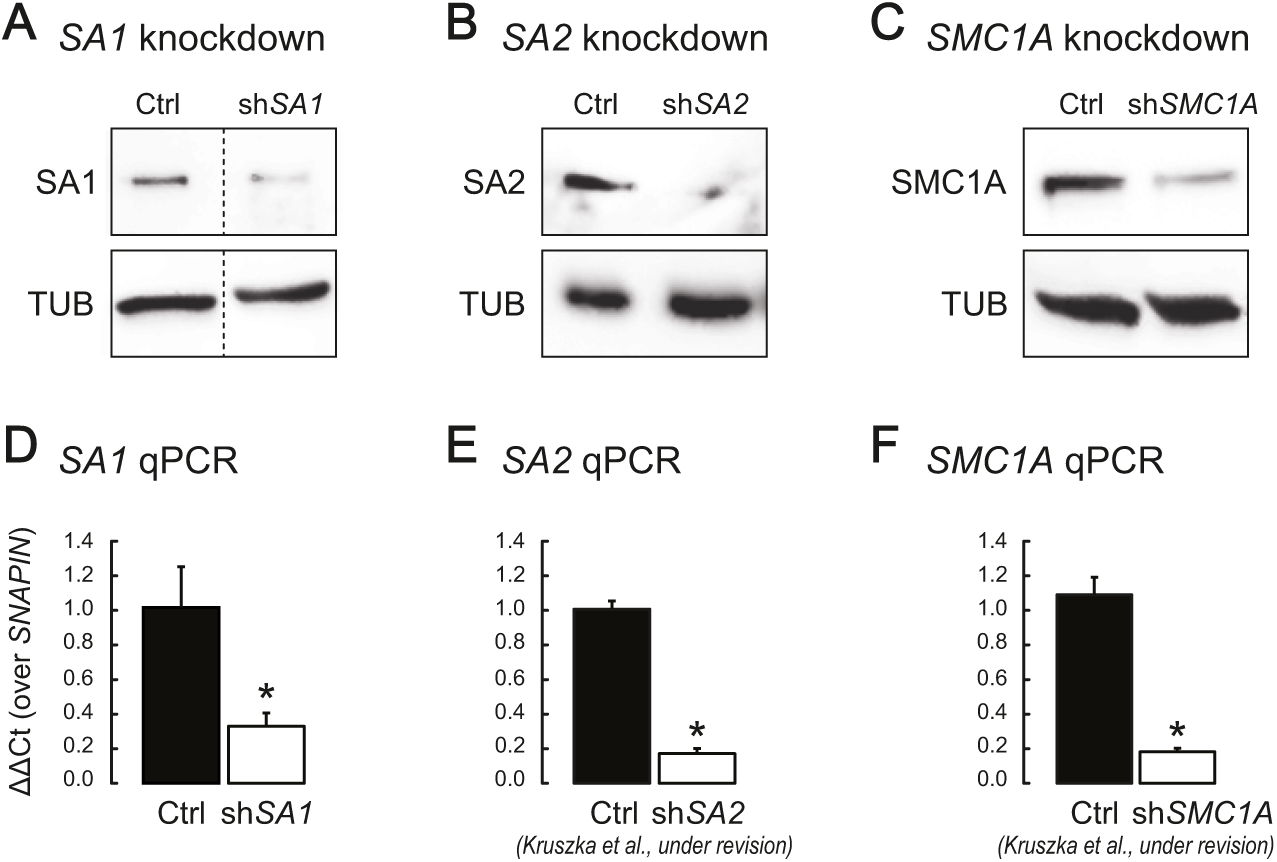
Depletion of cohesin from neural stem cells. **A**-**C**) Depletion of SA1, SA2 and SMC1A by siRNA-mediated depletion in neural stem cells was confirmed by Western blotting for SA1 (A), SA2 (B) and SMC1A (C). Tubulin (tub) was used as loading control. **D-F**) Depletion of SA1, SA2 and SMC1A by siRNA-mediated depletion in neural stem cells was confirmed by RT-PCR/qPCR analysis for the mRNA of *SA1, SA2 and SMC1A. SNAPIN* was used as housekeeping gene. Please note that some of these controls are also shown in Kruska et al. (Kruska et al., submitted).

**Figure S8.**
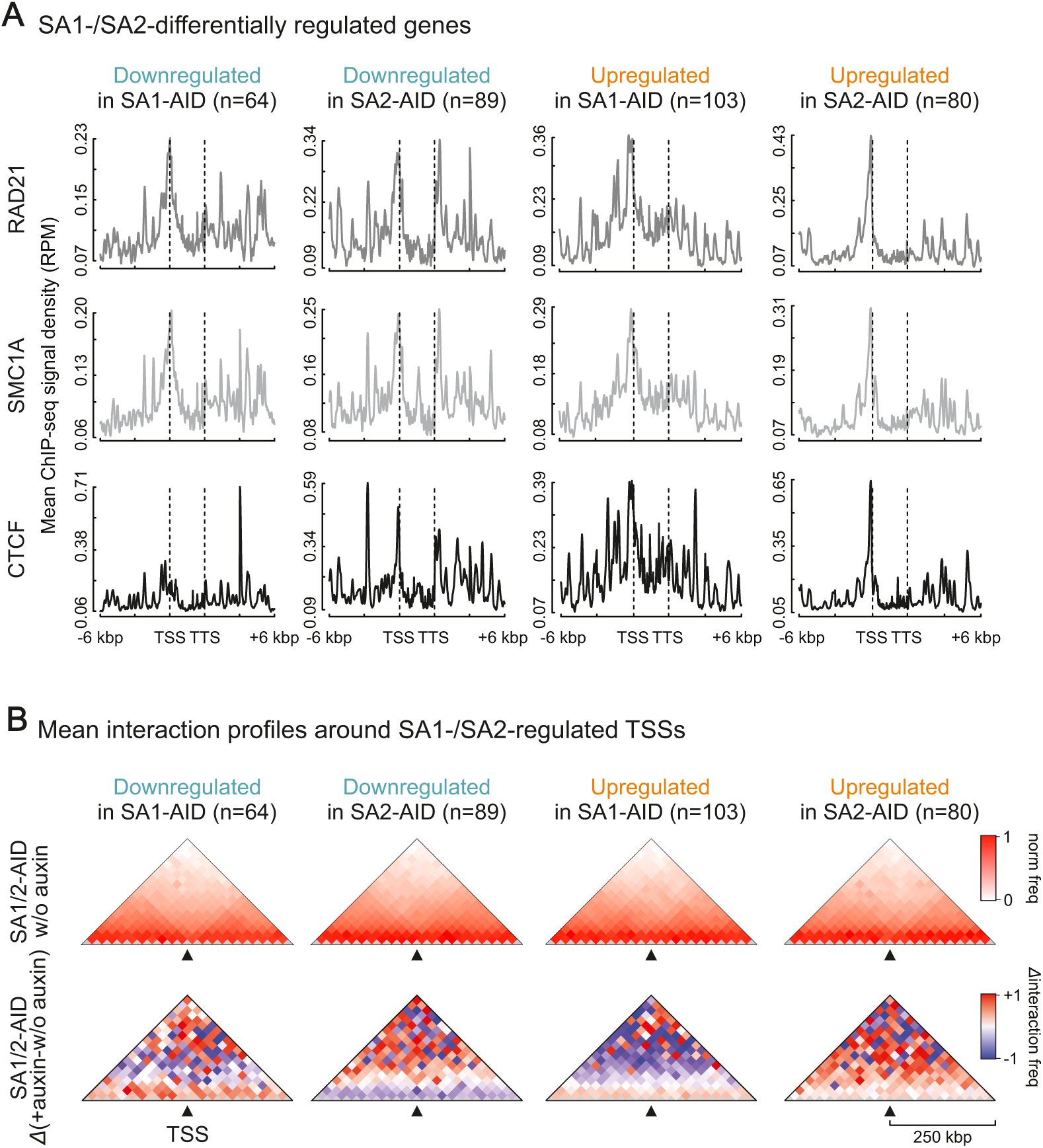
Analysis of gene subsets regulated after auxin-mediated depletion of SA1 or SA2. **A)** Mean RAD21, SMC1, and CTCF ChIP-seq signal was plotted along the bodies (±6 kbp) of genes specifically up- or down-regulated after depletion of SA1 or SA2. **B)** Insulation plots showing mean Hi-C signal from untreated maps (no auxin) around the TSSs of genes specifically up- or down-regulated after depletion of SA1 or SA2. For these same TSSs mean changes in Hi-C signal are also plotted. The maps are presented with a bin size of 25 kbp for the 20 bins around each TSS.

**Table S1.** Antibodies.

| **Target** | **Source** | **Use** | **Referred to** |
| --- | --- | --- | --- |
| α-GFP | #3611814460001, Roche | WB, IP/WB | Fig. 1 |
| α-EGFP | Homemade rabbit antibody | ChIP-seq, ChIP-qPCR, IF | Fig. 1, Fig. S1, Fig. 2, Fig. S2 |
| α-SA1 | Homemade rabbit antibody | One-Day ChIP Kit, Re-ChIP | Fig. S3 |
| α-SA1 | J.M. Peters | WB | Fig. 1 |
| α-SA1 | #A300-156A, Bethyl Laboratories | WB | Fig. S6 |
| α-SA2 | Homemade rabbit antibody | One-Day ChIP kit, Re-ChIP | Fig. S3 |
| α-SA2 | J.M. Peters | WB | Fig. 1 |
| α-SA2 | #A300-055A, Bethyl Laboratories | WB | Fig. S6 |
| α-SMC1A | Niels Galjart | IP/WB | Fig. 1 |
| α-SMC1A | #A300-055A, Bethyl Laboratories | WB | Fig. S6 |
| α-SMC3 | Homemade rabbit antibody | IP, ChIP-qPCR, Re-ChIP | Fig. 1, Fig. S3, Fig. S2 |
| Rabbit IgG | #C15410206, Diagenode | One-Day ChIP Kit, Re-ChIP | Fig. S3 |
| Rabbit IgG | #500-P00-500μg, Peprotech | IP, ChIP-qPCR | Fig. 1, Fig. 2, Fig. S2 |
| α-Tubulin | #T8328, Sigma | WB | Fig. 1, Fig. S6 |
| α-SA1 | Abcam ab4457 | dSTORM | Fig. 3 |
| α-SA2 | Bethyl A302-580A | dSTORM | Fig. 3 |
| α-CTCF | BD 612148 | dSTORM | Fig. 3, Fig. S3 |

**Table S2.**
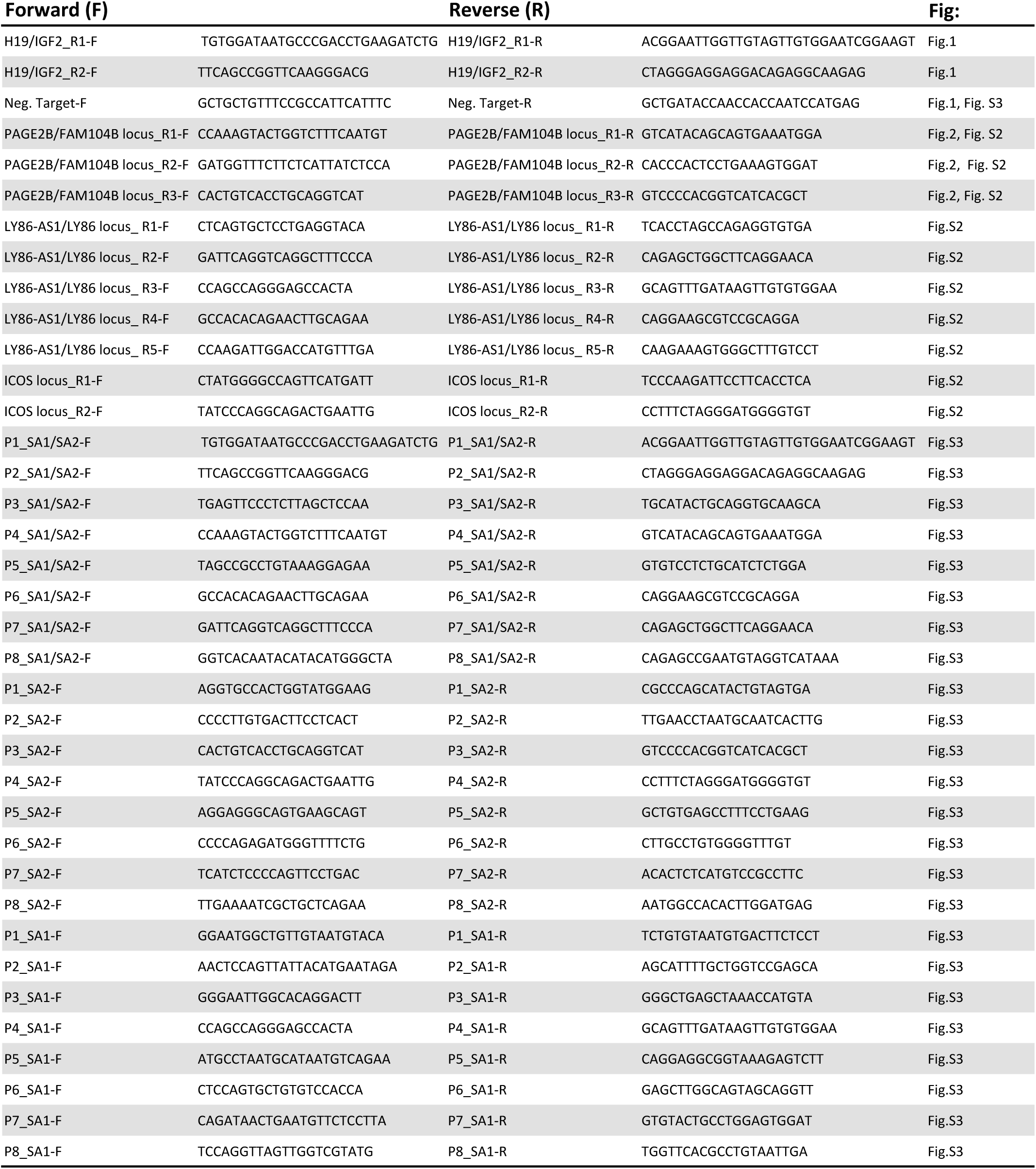
ChIP-qPCR primers.

**Table S3.**
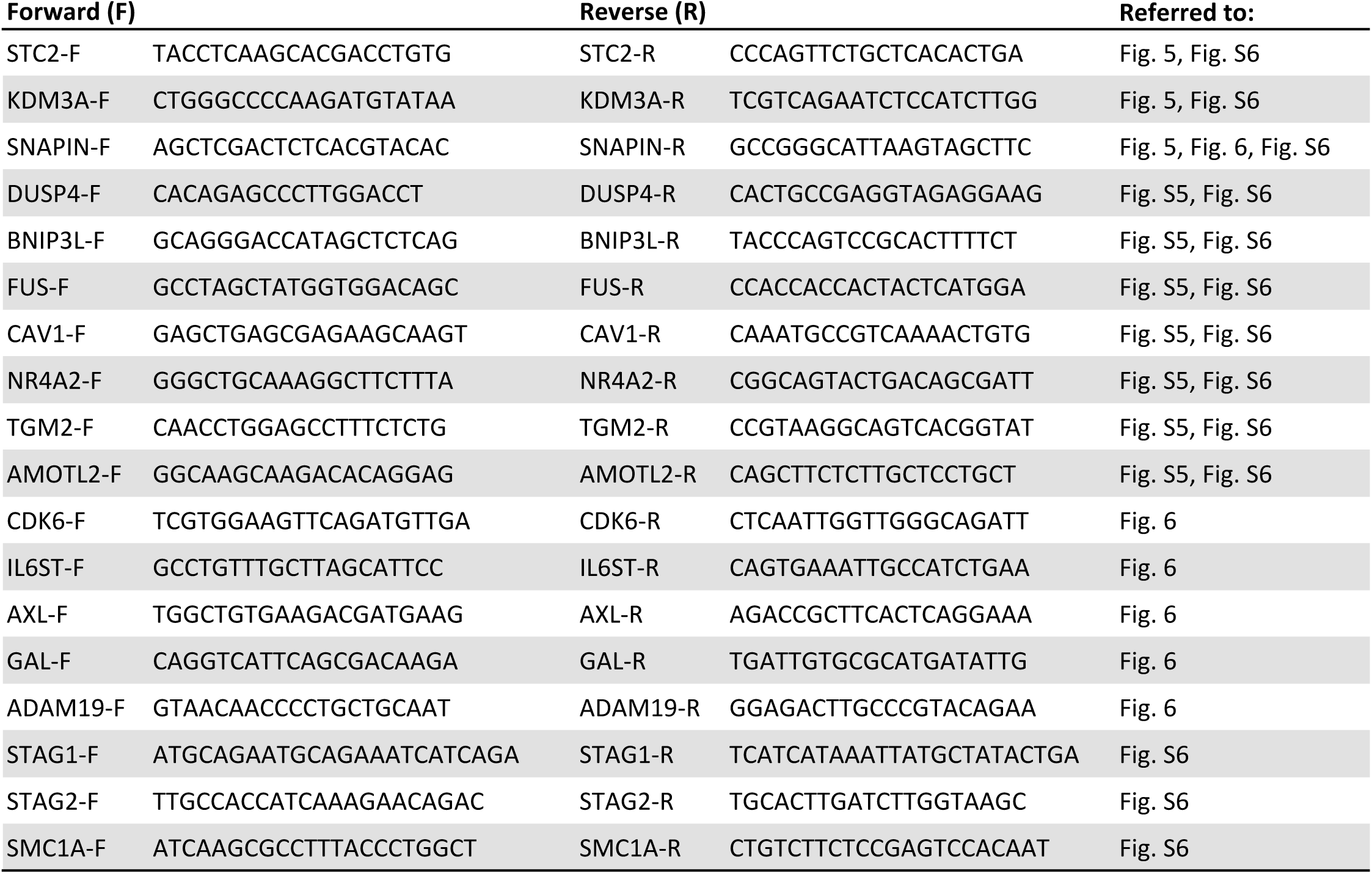
RT-qPCR primers.

**Table S4.** Differentially expressed genes after SA1 degradation (FDR < 0,05; log2 FC >0,6) (Genes annotated in bold red are genes that respond to auxin (Rao et al., Cell, 2017)

| Gene name | baseMean | log2FoldChange | pvalue | ENS Gene |
| --- | --- | --- | --- | --- |
| **STC2** | 1553,04307 | 0,429846247 | 3,63534E-05 | ENSG00000113739 |
| **AC007238.1** | 238,304234 | -0,223850444 | 0,03946788 | ENSG00000231043 |
| **AC013248.2** | 660,858416 | -0,232946589 | 0,035302666 | ENSG00000230897 |
| **AC015712.2** | 384,229879 | 0,488001244 | 1,32694E-05 | ENSG00000259583 |
| **AC026366.1** | 2413,79956 | -0,282700822 | 0,005533248 | ENSG00000240342 |
| **AC074033.1** | 253,40146 | -0,274798627 | 0,012235553 | ENSG00000213178 |
| **AC091167.1** | 140,066563 | -0,205942115 | 0,040986763 | ENSG00000225193 |
| **AC136632.1** | 457,307896 | -0,270154174 | 0,015947765 | ENSG00000218227 |
| **ADAM19** | 2129,74826 | 0,369498689 | 0,000382867 | ENSG00000135074 |
| **ADGRF1** | 120,730402 | 0,213286137 | 0,021349738 | ENSG00000153292 |
| **ADIRF-AS1** | 228,131195 | 0,309592231 | 0,004376332 | ENSG00000272734 |
| **ADORA2B** | 1282,01441 | 0,277295599 | 0,008307564 | ENSG00000170425 |
| **AHRR** | 650,355628 | 0,676510312 | 0,000000001 | ENSG00000063438 |
| **AKAP12** | 101645,75 | 0,247627334 | 0,005230597 | ENSG00000131016 |
| **AL009174.1** | 35,5021956 | -0,216570546 | 0,000969204 | ENSG00000227008 |
| **ALDH1A3** | 5522,92639 | 0,720718461 | 1,68579E-14 | ENSG00000184254 |
| **ANXA2P2** | 632,621689 | 0,22631971 | 0,005339164 | ENSG00000231991 |
| **AQP3** | 281,644987 | 0,254042966 | 0,021620495 | ENSG00000165272 |
| **ARL4D** | 288,551225 | 0,226624299 | 0,039159652 | ENSG00000175906 |
| **ARSG** | 260,940284 | 0,229507993 | 0,036831332 | ENSG00000141337 |
| **ATP11A** | 2243,80931 | 0,205816791 | 0,045838606 | ENSG00000068650 |
| **ATP5J** | 1763,87771 | -0,229295088 | 0,025785912 | ENSG00000154723 |
| **ATP9A** | 1392,45901 | 0,295727884 | 0,004933776 | ENSG00000054793 |
| **BHLHE40** | 1011,55291 | 0,269191215 | 0,012712364 | ENSG00000134107 |
| **BNIP3L** | 217,125089 | 0,291630964 | 0,00686662 | ENSG00000104765 |
| **BUB1B** | 1060,68306 | -0,224271385 | 0,03684633 | ENSG00000156970 |
| **C1orf116** | 237,116115 | 0,253547394 | 0,020110651 | ENSG00000182795 |
| **CAPN5** | 235,076964 | 0,225168047 | 0,038939534 | ENSG00000149260 |
| **CAV1** | 3774,08524 | -0,260436934 | 0,006888418 | ENSG00000105974 |
| **CBWD2** | 134,72796 | 0,203641035 | 0,036820121 | ENSG00000136682 |
| **CCNA2** | 807,479729 | -0,22992282 | 0,036447375 | ENSG00000145386 |
| **CD109** | 3053,91097 | 0,267953966 | 0,008764745 | ENSG00000156535 |
| **CD59** | 3938,52459 | 0,194168898 | 0,041691076 | ENSG00000085063 |
| **CDCA5** | 2176,02297 | -0,206312446 | 0,043041166 | ENSG00000146670 |
| **CDK1** | 1335,46082 | -0,235500243 | 0,027427664 | ENSG00000170312 |
| **CDK6** | 2650,48925 | 0,302153586 | 0,003364328 | ENSG00000105810 |
| **COX7C** | 2986,04004 | -0,240004655 | 0,013983887 | ENSG00000127184 |
| **CPA4** | 9395,53881 | 0,254305472 | 0,006808085 | ENSG00000128510 |
| **CREB3L2** | 1429,33861 | 0,21677937 | 0,037242233 | ENSG00000182158 |
| **CRY1** | 255,342622 | 0,226608196 | 0,038693403 | ENSG00000008405 |
| **CRY2** | 357,077891 | 0,235191803 | 0,035536866 | ENSG00000121671 |
| CTNNA1 | 9617,15462 | 0,221571227 | 0,015120528 | ENSG00000044115 |
| CTNNAL1 | 1821,80906 | -0,223999278 | 0,028853176 | ENSG00000119326 |
| CYP1A1 | 254,743646 | 0,923728579 | 3,52374E-17 | ENSG00000140465 |
| CYP1B1 | 103,303218 | 0,399898714 | 1,40768E-05 | ENSG00000138061 |
| DAGLB | 745,352963 | 0,226636061 | 0,039059288 | ENSG00000164535 |
| DDX39A | 1931,44745 | -0,214456134 | 0,038801073 | ENSG00000123136 |
| DHRS3 | 638,345269 | 0,291366772 | 0,008778444 | ENSG00000162496 |
| DUSP4 | 3871,47077 | 0,326901128 | 0,000637067 | ENSG00000120875 |
| EEF1A1 | 21324,5176 | 0,240528981 | 0,008221504 | ENSG00000156508 |
| EGFR | 2811,33006 | 0,24131266 | 0,016826998 | ENSG00000146648 |
| EREG | 6478,90214 | 0,283980333 | 0,002192839 | ENSG00000124882 |
| ERRFI1 | 10561,4611 | 0,214644763 | 0,022594264 | ENSG00000116285 |
| ESPL1 | 959,217933 | -0,234742213 | 0,029695875 | ENSG00000135476 |
| EXOSC9 | 1506,97899 | -0,21540063 | 0,037526859 | ENSG00000123737 |
| FANCG | 1139,51958 | -0,225326265 | 0,0370216 | ENSG00000221829 |
| FGF9 | 152,1991 | 0,28080702 | 0,00473005 | ENSG00000102678 |
| FUCA1 | 727,491446 | 0,331438148 | 0,00270783 | ENSG00000179163 |
| FUS | 6839,05855 | -0,325230148 | 0,000470144 | ENSG00000089280 |
| GAL | 939,272138 | -0,231878265 | 0,031874768 | ENSG00000069482 |
| GALNT5 | 3457,58473 | 0,277505524 | 0,006203653 | ENSG00000136542 |
| GAPDHP65 | 1026,85298 | -0,228233955 | 0,033629145 | ENSG00000235587 |
| GDA | 1632,64285 | 0,453899985 | 0,000015009 | ENSG00000119125 |
| GDF15 | 6593,3238 | 0,474156977 | 3,53918E-07 | ENSG00000130513 |
| GLI2 | 320,286431 | 0,239688862 | 0,029756323 | ENSG00000074047 |
| GNG12 | 1305,50407 | 0,208833409 | 0,049899282 | ENSG00000172380 |
| GPRC5A | 14470,1639 | 0,365431032 | 6,65725E-05 | ENSG00000013588 |
| H1FO | 7472,14175 | 0,279836254 | 0,002932592 | ENSG00000189060 |
| HAS3 | 2656,83242 | 0,908781683 | 2,6989E-20 | ENSG00000103044 |
| HBE1 | 1072,38293 | 0,244966296 | 0,022016039 | ENSG00000213931 |
| HECW2 | 137,684504 | 0,200967766 | 0,045804615 | ENSG00000138411 |
| HIPK2 | 2734,67007 | 0,358507202 | 0,00026019 | ENSG00000064393 |
| HMGB1 | 1479,49622 | -0,232494592 | 0,025781286 | ENSG00000189403 |
| HNRNPD | 5092,84814 | -0,207774607 | 0,028469751 | ENSG00000138668 |
| HPCAL1 | 1969,60217 | 0,442399889 | 1,42505E-05 | ENSG00000115756 |
| HSPB1 | 3767,12015 | -0,219196259 | 0,022266624 | ENSG00000106211 |
| IRF2BPL | 1436,32805 | 0,2556335 | 0,014416818 | ENSG00000119669 |
| ITGB5 | 1794,47167 | 0,222310303 | 0,029623772 | ENSG00000082781 |
| ITPR1 | 231,734143 | 0,220305252 | 0,042875571 | ENSG00000150995 |
| KHNYN | 1328,74673 | 0,209633771 | 0,045917647 | ENSG00000100441 |
| KIAA1683 | 229,589509 | 0,285833318 | 0,008592335 | ENSG00000130518 |
| KNTC1 | 1006,85372 | -0,250433006 | 0,020692262 | ENSG00000184445 |
| KRT15 | 917,824711 | 0,256939684 | 0,018128166 | ENSG00000171346 |
| LAMC2 | 1240,74127 | 0,262490347 | 0,012909499 | ENSG00000058085 |
| LDHAP5 | 299,224978 | -0,220346573 | 0,046475284 | ENSG00000213574 |
| LGALS3 | 1606,14482 | 0,464059179 | 6,86372E-06 | ENSG00000131981 |
| LIMA1 | 3038,72204 | 0,211729378 | 0,029455826 | ENSG00000050405 |
| LINC00511 | 596,648032 | 0,280642856 | 0,011624763 | ENSG00000227036 |
| **LMF2** | 981,906922 | -0,229311586 | 0,035153695 | ENSG00000100258 |
| **LPP** | 1197,9245 | 0,26713684 | 0,012377485 | ENSG00000145012 |
| **LTK** | 382,19436 | 0,231620684 | 0,037381964 | ENSG00000062524 |
| **MCM2** | 3691,6045 | -0,201265429 | 0,03694286 | ENSG00000073111 |
| **MCM3** | 6174,61263 | -0,185315573 | 0,049846859 | ENSG00000112118 |
| **MCM4** | 5619,3297 | -0,198772422 | 0,03301273 | ENSG00000104738 |
| **MED13L** | 5225,65175 | 0,194441332 | 0,038337255 | ENSG00000123066 |
| **MID1** | 3195,83089 | 0,397453862 | 4,62584E-05 | ENSG00000101871 |
| **MLPH** | 1927,1131 | 0,232293767 | 0,023922346 | ENSG00000115648 |
| **MT1G** | 313,839859 | 0,231708339 | 0,037280636 | ENSG00000125144 |
| **MT2A** | 4596,17198 | 0,324540696 | 0,000594269 | ENSG00000125148 |
| **MTCO3P12** | 15720,2806 | 0,24389268 | 0,006421804 | ENSG00000198744 |
| **MTUS1** | 1688,05962 | 0,228135481 | 0,026881805 | ENSG00000129422 |
| **MYEOV** | 3455,03849 | 0,224217226 | 0,022122473 | ENSG00000172927 |
| **MYOF** | 14243,616 | 0,201045953 | 0,027357519 | ENSG00000138119 |
| **NCAPG2** | 1490,01692 | -0,207076745 | 0,045682108 | ENSG00000146918 |
| **NDRG1** | 2969,30105 | 0,324724545 | 0,000860687 | ENSG00000104419 |
| **NGFR** | 325,583342 | 0,269895422 | 0,015441168 | ENSG00000064300 |
| **NPAS2** | 544,062208 | 0,265718547 | 0,017699227 | ENSG00000170485 |
| **NPTN** | 946,354944 | 0,214123093 | 0,047487458 | ENSG00000156642 |
| **NQO1** | 5373,86721 | 0,405755776 | 1,47446E-05 | ENSG00000181019 |
| **NUP85** | 2349,83045 | -0,214895386 | 0,032791494 | ENSG00000125450 |
| **NUSAP1** | 1374,80358 | -0,209239124 | 0,04660435 | ENSG00000137804 |
| **OR51B4** | 448,657817 | 0,308182647 | 0,005981971 | ENSG00000183251 |
| **OSGIN1** | 634,817481 | 0,254190389 | 0,022510548 | ENSG00000140961 |
| **OTUB2** | 368,217379 | 0,392204231 | 0,000454653 | ENSG00000089723 |
| **PAM** | 3606,50061 | 0,21638714 | 0,024117499 | ENSG00000145730 |
| **PCLAF** | 559,871692 | -0,22104183 | 0,047662877 | ENSG00000166803 |
| **PDLIM5** | 2830,15734 | 0,195829374 | 0,045259246 | ENSG00000163110 |
| **PER2** | 163,218515 | 0,261224339 | 0,011464355 | ENSG00000132326 |
| **PNN** | 4450,05548 | -0,202040971 | 0,033390779 | ENSG00000100941 |
| **POLE** | 3394,58564 | -0,197166136 | 0,045192495 | ENSG00000177084 |
| **PPIAP29** | 75,9083929 | -0,485157062 | 0,000000007 | ENSG00000214975 |
| **RABGGTB** | 2013,42251 | -0,216703188 | 0,03192614 | ENSG00000137955 |
| **RANBP1** | 1852,04814 | -0,255038216 | 0,012023616 | ENSG00000099901 |
| **RNF128** | 168,584917 | 0,239852253 | 0,021259165 | ENSG00000133135 |
| **RPL34** | 4460,86115 | -0,201320036 | 0,035633711 | ENSG00000109475 |
| **RPL5P4** | 42,4816015 | -0,227343704 | 0,00065862 | ENSG00000229994 |
| **RPS23** | 4019,23236 | -0,227336628 | 0,023767614 | ENSG00000186468 |
| **RPS27AP16** | 3534,72097 | -0,231096045 | 0,019727652 | ENSG00000224631 |
| **RPS29** | 4474,3423 | -0,259061353 | 0,008158282 | ENSG00000213741 |
| **RPS6KA2** | 226,295931 | 0,304341601 | 0,003667454 | ENSG00000071242 |
| **RPSA** | 4154,11797 | -0,277403214 | 0,003734191 |  |
| **RRM1** | 2456,79125 | -0,284560695 | 0,004156358 | ENSG00000167325 |
| **SECTM1** | 529,907198 | 0,369123808 | 0,000955381 | ENSG00000141574 |
| **SH3BP4** | 1568,61242 | 0,24950967 | 0,018066042 | ENSG00000130147 |
| **SH3KBP1** | 1813,24675 | 0,213684426 | 0,038497203 | ENSG00000147010 |
| SLC16A6 | 121,126739 | 0,201147792 | 0,03997311 | ENSG00000108932 |
| SLC2A1 | 6167,59549 | 0,294675487 | 0,001584407 | ENSG00000117394 |
| SLC37A2 | 77,0629637 | 0,219361769 | 0,011711534 | ENSG00000134955 |
| SLC39A11 | 726,747305 | 0,226651691 | 0,040595181 | ENSG00000133195 |
| SLC3A2 | 8249,46266 | 0,310292274 | 0,000805144 | ENSG00000168003 |
| SLC7A5 | 13624,4039 | 0,323735834 | 0,000334664 | ENSG00000103257 |
| SMAD3 | 2625,60913 | 0,345543981 | 0,000506292 | ENSG00000166949 |
| SMIM14 | 257,261789 | 0,215990339 | 0,049473011 | ENSG00000163683 |
| SNHG1 | 5387,94725 | -0,199260323 | 0,033787959 | ENSG00000255717 |
| SNHG25 | 150,436777 | -0,217940219 | 0,031762084 | ENSG00000266402 |
| SNHG9 | 534,118578 | -0,227590459 | 0,042317473 | ENSG00000255198 |
| SNRPG | 605,311887 | -0,220747636 | 0,047073183 | ENSG00000143977 |
| SOX9 | 2149,535 | 0,250003085 | 0,014875191 | ENSG00000125398 |
| SRSF1 | 5638,93868 | -0,19119989 | 0,041665942 | ENSG00000136450 |
| SSH1 | 4842,92711 | 0,306130273 | 0,001194932 | ENSG00000084112 |
| SUMO1P3 | 62,8002729 | -0,21484975 | 0,003179657 | ENSG00000235082 |
| TEF | 236,536659 | 0,305396122 | 0,004993677 | ENSG00000167074 |
| TGFA | 1743,24663 | 0,239667746 | 0,018916938 | ENSG00000163235 |
| THOC3 | 355,803421 | -0,248067427 | 0,026585883 | ENSG00000051596 |
| TIMM23B | 27,9068186 | -0,145820157 | 0,013440218 | ENSG00000204152 |
| TIPARP | 864,015609 | 0,505235741 | 4,89519E-06 | ENSG00000163659 |
| TLDC1 | 1287,93974 | 0,31058647 | 0,003163448 | ENSG00000140950 |
| TMEM2 | 909,943339 | 0,217213992 | 0,045012772 | ENSG00000135048 |
| TMEM200A | 428,63253 | 0,260951785 | 0,01995854 | ENSG00000164484 |
| TMEM221 | 75,4495689 | 0,176376876 | 0,041593652 | ENSG00000188051 |
| TOMM40 | 2880,31969 | -0,204492532 | 0,037347492 | ENSG00000130204 |
| TPM2 | 7085,40699 | -0,212072887 | 0,024327693 | ENSG00000198467 |
| TRIM2 | 665,133329 | 0,341136626 | 0,002064531 | ENSG00000109654 |
| TRIOBP | 808,807088 | 0,219404386 | 0,044688258 | ENSG00000100106 |
| TSC22D1 | 6699,60893 | 0,35883103 | 0,000324115 | ENSG00000102804 |
| TSPAN18 | 339,119595 | 0,21846273 | 0,046744187 | ENSG00000157570 |
| TUBA1B | 2116,57294 | -0,239828003 | 0,01837716 | ENSG00000123416 |
| TUBA1C | 1542,3865 | -0,203094852 | 0,049369336 | ENSG00000167553 |
| TYMS | 1427,83029 | -0,231660138 | 0,026673775 | ENSG00000176890 |
| UBE2C | 926,394729 | -0,248847799 | 0,022068283 | ENSG00000175063 |
| UNC13A | 3245,49718 | 0,194700436 | 0,0442065 | ENSG00000130477 |
| USP39 | 1923,84998 | -0,218147507 | 0,031100452 | ENSG00000168883 |
| WDR34 | 1471,11155 | -0,22324974 | 0,034859573 | ENSG00000119333 |
| WNT16 | 475,172545 | 0,290349912 | 0,009566714 | ENSG00000002745 |
| WNT9A | 672,65464 | 0,260004 | 0,018641762 | ENSG00000143816 |
| ZFP36L1 | 2953,26356 | 0,296502364 | 0,002346183 | ENSG00000185650 |
| ZWINT | 1913,28877 | -0,245259449 | 0,015680565 | ENSG00000122952 |

**Table S5.** Differentially expressed genes after SA2 degradation (FDR < 0,05; log2 FC >0,6) (Genes annotated in bold red are genes that respond to auxin (Rao et al., Cell, 2017)

| Gene name | baseMean | log2FoldChange | pvalue | ENS Gene |
| --- | --- | --- | --- | --- |
| AC015712.2 | 384,8367021 | 0,343536448 | 0,003846459 | ENSG00000259583 |
| AC026254.2 | 122,2613771 | 0,501084233 | 1,38642E-06 | ENSG00000239223 |
| AC040162.1 | 76,25874216 | -0,186694117 | 0,045375069 | ENSG00000132382 |
| AC073621.1 | 3563,864194 | -0,234192151 | 0,029087202 | ENSG00000178896 |
| AC106795.1 | 85,64286778 | -0,244098067 | 0,012887103 | ENSG00000183087 |
| ACTN4 | 12024,8453 | -0,226798952 | 0,027817609 | ENSG00000013375 |
| ADD3 | 663,1215888 | 0,27080804 | 0,021368274 | ENSG00000067167 |
| AGRN | 2281,53145 | -0,237771088 | 0,040266694 | ENSG00000164362 |
| **AHRR** | 411,5171477 | 0,613320808 | 0,00000025 | ENSG00000063438 |
| AL049714.1 | 43,27995222 | 0,277092103 | 0,000243308 | ENSG00000115548 |
| AL109741.2 | 28,09865026 | -0,156925872 | 0,021503845 | ENSG00000150961 |
| AL358472.1 | 15,89799953 | 0,11332353 | 0,032748936 | ENSG00000110047 |
| **ALDH1A3** | 4487,604962 | 0,372165531 | 0,000787307 | ENSG00000184254 |
| ALYREF | 2637,710198 | -0,223946285 | 0,046348333 | ENSG00000099624 |
| AMOTL2 | 1290,77886 | -0,380629925 | 0,001156461 | ENSG00000187961 |
| ANKRD12 | 279,3924239 | 0,273362385 | 0,018955028 | ENSG00000099795 |
| AQP3 | 204,3853501 | 0,2734884 | 0,014623327 | ENSG00000165272 |
| ARHGEF37 | 332,5581374 | 0,304375561 | 0,010230253 | ENSG00000114019 |
| ASPSCR1 | 728,5663998 | -0,238282752 | 0,043706312 | ENSG00000025770 |
| ATAD3B | 1207,909712 | -0,247737829 | 0,031501388 | ENSG00000188157 |
| ATP5D | 1232,25509 | -0,258024608 | 0,028864409 | ENSG00000198380 |
| AURKAIP1 | 1479,549143 | -0,226988725 | 0,047635975 | ENSG00000127586 |
| AURKB | 748,1378103 | -0,278142864 | 0,018907323 | ENSG00000130702 |
| AXL | 1925,996356 | -0,259044996 | 0,020607361 | ENSG00000176018 |
| CADPS2 | 267,1602114 | 0,28054561 | 0,016340993 | ENSG00000063660 |
| CAPN15 | 646,0101997 | -0,235655075 | 0,047522829 | ENSG00000231416 |
| CAV1 | 4663,658632 | -0,280171021 | 0,008840816 | ENSG00000105974 |
| CAVIN1 | 5760,066965 | -0,264975019 | 0,014541408 | ENSG00000034152 |
| CCDC186 | 612,6470283 | 0,262164935 | 0,026457333 | ENSG00000155304 |
| CD109 | 3137,602527 | 0,341677968 | 0,002433663 | ENSG00000156535 |
| CDK6 | 3741,178881 | 0,226896153 | 0,033585778 | ENSG00000105810 |
| CEP170B | 2421,745841 | -0,235735064 | 0,040818566 | ENSG00000142627 |
| CGN | 706,0781858 | -0,270723202 | 0,022196092 | ENSG00000125912 |
| CHTF18 | 1103,064818 | -0,25381691 | 0,029802969 | ENSG00000144369 |
| CKB | 6217,640042 | -0,216102872 | 0,043918714 | ENSG00000186472 |
| CLUH | 4172,653194 | -0,264729579 | 0,016553663 | ENSG00000166165 |
| COL18A1 | 1141,378117 | -0,233037828 | 0,04801685 | ENSG00000204634 |
| COL9A3 | 854,5258926 | -0,277186141 | 0,017783767 | ENSG00000183684 |
| CREBRF | 111,1846609 | 0,210218014 | 0,044523028 | ENSG00000087077 |
| CYC1 | 4047,304583 | -0,233313079 | 0,028975619 | ENSG00000196878 |
| **CYP1A1** | 207,8273184 | 0,650896061 | 1,4891E-08 | ENSG00000140465 |
| **CYP1B1** | 135,8637742 | 0,426014603 | 8,67938E-05 | ENSG00000138061 |
| DDX51 | 982,8234669 | -0,27168034 | 0,018891634 | ENSG00000071794 |
| **DHRS3** | 489,7517385 | 0,275410525 | 0,020394665 | ENSG00000162496 |
| DISP1 | 215,6501615 | 0,230447702 | 0,04592773 | ENSG00000140451 |
| DPP7 | 2305,715134 | -0,239246214 | 0,035816302 | ENSG00000186480 |
| DUS3L | 610,2479378 | -0,259084161 | 0,028564112 | ENSG00000118515 |
| EBNA1BP2 | 2612,082088 | -0,224574619 | 0,038584023 | ENSG00000179091 |
| EEF1D | 6776,457067 | -0,243473236 | 0,025370621 | ENSG00000112144 |
| EGFR | 2044,138496 | 0,250170814 | 0,023118884 | ENSG00000146648 |
| EGR1 | 2945,432344 | 0,256977408 | 0,019345745 | ENSG00000006459 |
| EHD1 | 1066,519573 | -0,266773626 | 0,023215396 | ENSG00000164463 |
| EPHA2 | 5812,818836 | -0,22277592 | 0,033675944 | ENSG00000130402 |
| EXOSC4 | 750,0340241 | -0,252137566 | 0,032286558 | ENSG00000162006 |
| F3 | 1074,564505 | -0,294334615 | 0,010257585 | ENSG00000184990 |
| FAM102B | 301,0361861 | 0,287378005 | 0,0150974 | ENSG00000106366 |
| FAM171B | 383,5676569 | 0,24258572 | 0,041435565 | ENSG00000132361 |
| FGF9 | 204,9016955 | 0,287919095 | 0,012124657 | ENSG00000102678 |
| FGF9 | 980,5573876 | 0,294500898 | 0,010600599 | ENSG00000179163 |
| FGFBP1 | 98,86684065 | 0,201133171 | 0,048073726 | ENSG00000183751 |
| FLNC | 5341,900359 | -0,310100082 | 0,004829358 | ENSG00000137440 |
| FOS | 2082,444442 | 0,401701411 | 0,000419404 | ENSG00000153234 |
| GAL | 788,2632445 | -0,268459136 | 0,021611255 | ENSG00000069482 |
| GAS6 | 1097,599255 | -0,245019481 | 0,037790506 | ENSG00000138593 |
| GDA | 1927,082242 | 0,45294776 | 4,22538E-05 | ENSG00000119125 |
| GDF15 | 6345,725409 | 0,219630837 | 0,039037869 | ENSG00000130513 |
| GFPT1 | 1750,15019 | 0,227677668 | 0,041725319 | ENSG00000228399 |
| GJB3 | 716,7578616 | -0,277188612 | 0,019634363 | ENSG00000086544 |
| GPC1 | 2261,663728 | -0,289576574 | 0,008359392 | ENSG00000175756 |
| GULP1 | 671,7906819 | 0,256602572 | 0,02927724 | ENSG00000177469 |
| H1FO | 5305,533346 | 0,242777375 | 0,025572273 | ENSG00000189060 |
| H2AFX | 1239,53394 | -0,236993874 | 0,041817481 | ENSG00000127666 |
| HAS3 | 1112,561461 | 0,515914842 | 8,25834E-06 | ENSG00000103044 |
| HAT1 | 1012,52403 | 0,269161528 | 0,02013686 | ENSG00000177700 |
| HIPK2 | 1882,110089 | 0,253191977 | 0,023429218 | ENSG00000064393 |
| HLTF | 981,9837767 | 0,231061377 | 0,045918587 | ENSG00000128708 |
| HPCAL1 | 1088,530876 | 0,252071067 | 0,028906441 | ENSG00000115756 |
| HSPA13 | 715,5878159 | 0,288654747 | 0,014045111 | ENSG00000175793 |
| ICK | 295,1385933 | 0,246247733 | 0,037120649 | ENSG00000135480 |
| IDH1 | 1200,981012 | 0,228791114 | 0,044380748 | ENSG00000167601 |
| IL6ST | 477,3367111 | 0,314802296 | 0,008140836 | ENSG00000244535 |
| INF2 | 1492,338932 | -0,238688038 | 0,03862448 | ENSG00000176978 |
| INSIG1 | 728,3856007 | 0,245477444 | 0,036286497 | ENSG00000165684 |
| INTS8 | 962,2908645 | 0,238296553 | 0,041197772 | ENSG00000196756 |
| ISYNA1 | 2709,390161 | -0,217193478 | 0,048261711 | ENSG00000167755 |
| ITPKC | 538,341544 | -0,2368878 | 0,046388887 | ENSG00000128626 |
| ITPR1 | 324,2172668 | 0,30954599 | 0,009016786 | ENSG00000150995 |
| ITPR2 | 345,9874576 | 0,276300912 | 0,019854484 | ENSG00000117525 |
| KDM3A | 1034,265164 | 0,299562994 | 0,009732671 | ENSG00000198918 |
| KDM7A | 452,5775543 | 0,291407766 | 0,01423637 | ENSG00000152558 |
| KIAA1683 | 176,1416106 | 0,262822415 | 0,020026857 | ENSG00000130518 |
| KLHL17 | 1082,713 | -0,230375834 | 0,048685769 | ENSG00000166166 |
| KLK6 | 673,1270016 | -0,263709391 | 0,025402083 | ENSG00000169696 |
| KRT7 | 289,5259415 | -0,280506998 | 0,015448656 | ENSG00000177707 |
| KRT8 | 38619,73064 | -0,216112054 | 0,040117001 | ENSG00000109680 |
| LAMA5 | 6063,177906 | -0,209582077 | 0,046868698 | ENSG00000070814 |
| LAMB3 | 2022,157194 | -0,236393924 | 0,032095645 | ENSG00000114767 |
| LGALS3 | 1149,289095 | 0,238376356 | 0,038969489 | ENSG00000131981 |
| LINC00511 | 593,406884 | 0,249583433 | 0,03522631 | ENSG00000227036 |
| LPP | 1315,656376 | 0,254391796 | 0,024673182 | ENSG00000145012 |
| LYSMD3 | 353,8116475 | 0,240631739 | 0,042833246 | ENSG00000006062 |
| MAN2A1 | 827,493203 | 0,261093879 | 0,026215756 | ENSG00000125731 |
| MAP2K3 | 2742,910921 | -0,227234177 | 0,04415226 | ENSG00000169750 |
| MAP3K14 | 366,7247578 | -0,280866518 | 0,017959409 | ENSG00000099994 |
| MID1 | 3453,768669 | 0,260377991 | 0,016585339 | ENSG00000101871 |
| MRPL45P2 | 171,8077608 | -0,27296757 | 0,014219748 | ENSG00000154309 |
| MRPS12 | 1444,475199 | -0,245245492 | 0,033917484 | ENSG00000117395 |
| MSLNL | 122,3519433 | -0,216989116 | 0,041262123 | ENSG00000104529 |
| MYBBP1A | 2937,990729 | -0,247572379 | 0,024626946 | ENSG00000170421 |
| MYOF | 15526,78597 | 0,305241035 | 0,003946188 | ENSG00000138119 |
| NCAPD2 | 3273,818249 | -0,225217483 | 0,03766847 | ENSG00000215030 |
| NCAPH2 | 1149,808233 | -0,246719737 | 0,035197268 | ENSG00000160551 |
| NCLN | 2895,799697 | -0,226353639 | 0,045575982 | ENSG00000144366 |
| NCOA7 | 254,7815986 | 0,232012485 | 0,046950973 | ENSG00000123104 |
| NDRG1 | 3413,449341 | 0,238381116 | 0,027756228 | ENSG00000104419 |
| NDUFB7 | 862,7152708 | -0,297873643 | 0,010896228 | ENSG00000168268 |
| NECTIN3 | 2423,464888 | 0,216455261 | 0,047070608 | ENSG00000143375 |
| NQO1 | 12579,85348 | 0,413677653 | 5,88011E-05 | ENSG00000181019 |
| NR1D2 | 819,7187178 | 0,260829779 | 0,026441788 | ENSG00000228782 |
| NR4A2 | 1977,967525 | 0,438077753 | 0,000117802 | ENSG00000198959 |
| NT5DC2 | 2015,58192 | -0,218213362 | 0,047792317 | ENSG00000112893 |
| OR51B4 | 531,1102715 | 0,310343401 | 0,008919742 | ENSG00000183251 |
| PCLO | 296,3343675 | 0,263676181 | 0,02527627 | ENSG00000230837 |
| PER2 | 156,1994998 | 0,230160941 | 0,038591881 | ENSG00000132326 |
| PGM3 | 1014,781984 | 0,25283608 | 0,029655782 | ENSG00000162636 |
| PIDD1 | 1080,740111 | -0,310881331 | 0,008269146 | ENSG00000105655 |
| PIF1 | 252,3583 | -0,266261395 | 0,020524855 | ENSG00000138413 |
| PJA2 | 993,1522887 | 0,267140243 | 0,021774063 | ENSG00000170089 |
| POLR2L | 1961,234912 | -0,286968381 | 0,01221301 | ENSG00000111961 |
| PTPN12 | 1412,660759 | 0,283475013 | 0,012992089 | ENSG00000177595 |
| PTPN23 | 1226,787425 | -0,247758442 | 0,029906623 | ENSG00000099814 |
| RAC3 | 1656,880499 | -0,248901749 | 0,030420278 | ENSG00000197747 |
| RBM14 | 1727,665015 | -0,222285957 | 0,047258258 | ENSG00000165813 |
| RHOT2 | 1005,553894 | -0,231050831 | 0,048861633 | ENSG00000160072 |
| RPL31P2 | 334,2277484 | 0,292607259 | 0,013579113 | ENSG00000128591 |
| RPL39 | 746,4029417 | 0,267099558 | 0,002959655 | ENSG00000127947 |
| RRP9 | 1254,078217 | -0,244004533 | 0,033683822 | ENSG00000010292 |
| S100A10 | 2877,881437 | -0,220834091 | 0,041095855 | ENSG00000174738 |
| SASH1 | 396,5967944 | 0,235952815 | 0,047307752 | ENSG00000178999 |
| SAT1 | 2914,622724 | 0,218978105 | 0,043373425 | ENSG00000151229 |
| SCRIB | 3168,339067 | -0,280010073 | 0,013272722 | ENSG00000111912 |
| SEC24D | 1025,079561 | 0,263857273 | 0,023441518 | ENSG00000180900 |
| SECISBP2L | 912,3232579 | 0,24173657 | 0,03818116 | ENSG00000092758 |
| SERPINE1 | 1480,843364 | -0,2997523 | 0,007840159 | ENSG00000182871 |
| **SESN3** | 123,0361905 | 0,221517343 | 0,03664887 | ENSG00000198961 |
| **SFN** | 9214,206238 | -0,28127354 | 0,007010089 | ENSG00000239306 |
| **SGK1** | 234,7396969 | 0,233243332 | 0,043221827 | ENSG00000120738 |
| **SH2D3A** | 1053,416342 | -0,288006267 | 0,01417384 | ENSG00000103326 |
| **SIVA1** | 1159,492457 | -0,230004723 | 0,047783612 | ENSG00000106688 |
| **SLC1A1** | 178,5993237 | 0,237069543 | 0,030704792 | ENSG00000185163 |
| **SLC2A13** | 113,6254193 | 0,237956624 | 0,021975258 | ENSG00000176014 |
| **SLC3A2** | 9305,865437 | 0,238024255 | 0,02207227 | ENSG00000168003 |
| **SMCHD1** | 1509,041118 | 0,276336347 | 0,018860998 | ENSG00000183111 |
| **SNAPC4** | 1294,6233 | -0,275097406 | 0,015865243 | ENSG00000116266 |
| **SNHG17** | 1099,592109 | -0,270046867 | 0,01837095 | ENSG00000261884 |
| **STXBP3** | 477,716723 | 0,237319308 | 0,04601972 | ENSG00000101596 |
| **SUSD2** | 222,2641519 | -0,227153705 | 0,047190006 | ENSG00000076201 |
| **TAOK1** | 2301,836082 | 0,232506087 | 0,0400913 | ENSG00000188910 |
| **TBC1D19** | 99,03038513 | 0,227758958 | 0,025353386 | ENSG00000148700 |
| **TBC1D8** | 1122,634323 | 0,24498009 | 0,032083011 | ENSG00000081803 |
| **TBL3** | 1213,473172 | -0,262769071 | 0,023033875 | ENSG00000169764 |
| **TCOF1** | 3392,883745 | -0,243251119 | 0,02499493 | ENSG00000149212 |
| **TERT** | 240,4919471 | -0,245402926 | 0,034894581 | ENSG00000203485 |
| **TGM2** | 1809,073465 | -0,453533862 | 7,00679E-05 | ENSG00000140983 |
| **TICAM1** | 669,0536322 | -0,249960408 | 0,034604723 | ENSG00000164941 |
| **TIPARP** | 957,0454751 | 0,429603366 | 0,000219342 | ENSG00000163659 |
| **TMEM123** | 1984,87583 | 0,309688406 | 0,00524705 | ENSG00000170345 |
| **TOMM40** | 2500,045643 | -0,239441188 | 0,031464074 | ENSG00000130204 |
| **TPM2** | 6939,431946 | -0,245223449 | 0,024744808 | ENSG00000198467 |
| **TRAM1** | 1878,899316 | 0,285449218 | 0,011926022 | ENSG00000134352 |
| **TRIM2** | 968,8913394 | 0,486572969 | 2,84237E-05 | ENSG00000109654 |
| **TRIP6** | 2179,832996 | -0,258026511 | 0,022953363 | ENSG00000188486 |
| **TRMT61A** | 596,5524084 | -0,256795292 | 0,030794077 | ENSG00000188229 |
| **TSC22D1** | 4217,883338 | 0,244379772 | 0,025030908 | ENSG00000102804 |
| **TUBB4B** | 7007,135157 | -0,223535862 | 0,04149241 | ENSG00000141994 |
| **TUBB6** | 2399,494472 | -0,263347078 | 0,018488506 | ENSG00000130066 |
| **UBE2C** | 1080,816529 | -0,303345574 | 0,009324256 | ENSG00000175063 |
| **UGP2** | 920,1915504 | 0,233723922 | 0,043330412 | ENSG00000101745 |

**Table S6.**
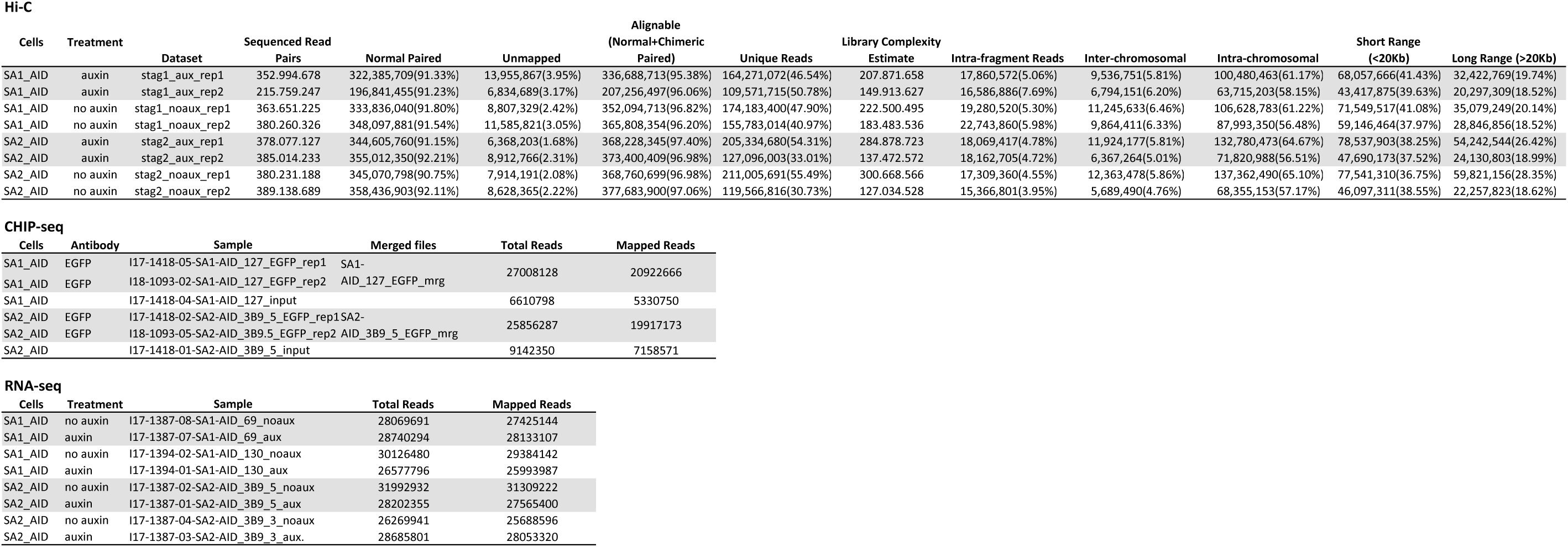
Sequencing data information and statistics.

**Table S7.** Peak statistics.

| Feature |  | Characteristics |  | Distribution of peaks (%) |  |  |
| --- | --- | --- | --- | --- | --- | --- |
|  |  |  |  | Common peaks | SA1-only peaks | SA2-only peaks |
| Promoter | active | TSS | H3K4me3 H3K27Ac | 9,5 | 8,0 | 32,6 |
|  | inactive | TSS | H3K4me3 | 0,3 | 0,2 | 7,0 |
| Enhancer | active | no promoter | H3K4me1 H3K27Ac | 15,4 | 14,9 | 12,1 |
|  | inactive | no promoter | H3K4me1 | 10,7 | 9,2 | 6,9 |
| Others |  |  |  | 64,1 | 67,7 | 41,4 |

**Table S8.** Published datasets used in this study.

| **Data Type** | **Reference** | **Website** |
| --- | --- | --- |
| RAD21 ChIP-seq | Rao et al., 2017 | https://www.ncbi.nlm.nih.gov/geo/query/acc.cgi?acc=GSM2809609 |
| CTCF ChIP-seq | Rao et al., 2017 | https://www.ncbi.nlm.nih.gov/geo/query/acc.cgi?acc=GSM2809615 |
| Input ChIP-seq for RAD21 and CTCF | Rao et al., 2017 | https://www.ncbi.nlm.nih.gov/geo/query/acc.cgi?acc=GSM3242976 |
| USF1 ChIP-seq | Davis CA et al., 2018 | https://www.encodeproject.org/experiments/ENCSR000BVK/ |
| FOSL1 ChIP-seq | Davis CA et al., 2018 | https://www.encodeproject.org/experiments/ENCSR000BTE/ |
| CEBPB ChIP-seq | Davis CA et al., 2018 | https://www.encodeproject.org/experiments/ENCSR000BSD/ |
| Input ChIP-seq for USF1, FOSL1 and CEBPB | Davis CA et al., 2018 | https://www.encodeproject.org/experiments/ENCSR000BMK/ |
| TCF7L2 ChIP-seq | Davis CA et al., 2018 | https://www.encodeproject.org/experiments/ENCSR000EUV/ |
| Input ChIP-seq for TCF7L2 | Davis CA et al., 2018 | https://www.encodeproject.org/experiments/ENCSR000EUX/ |
| H2A.Z ChIP-seq | Davis CA et al., 2018 | https://www.encodeproject.org/experiments/ENCSR227XNT/ |
| H3K4me1 ChIP-seq | Davis CA et al., 2018 | https://www.encodeproject.org/experiments/ENCSR161MXP/ |
| Input ChIP-seq for H2A.Z and HeK4me1 | Davis CA et al., 2018 | https://www.encodeproject.org/experiments/ENCSR198WIH/ |
| H3K4me3 ChIP-seq | Davis CA et al., 2018 | https://www.encodeproject.org/experiments/ENCSR000DTQ/ |
| Input ChIP-seq for H3K4me3 | Davis CA et al., 2018 | https://www.encodeproject.org/experiments/ENCSR000DTP/ |
| Rao et al., 2017 | Rao SSP, Huang SC, Glenn St Hilaire B, Engreitz JM et al.<br>Cohesin Loss Eliminates All Loop Domains. Cell 2017 Oct 5;171(2):305-320.e24. PMID: 28985562 |  |
| Davis CA et al., 2018 | Davis CA et al. The Encyclopedia of DNA elements (ENCODE): data portal update.<br>Nucleic Acids Res 2018. Jan 4;46(D1):D794-D801. PMID: 29126249 |  |

**Table S9.** Sequencing data information and statistics.

| **Software** | **Reference** | **Website** |
| --- | --- | --- |
| JUICER | Durand et al., 2016 | https://github.com/theaidenlab/juicer/wiki |
| GMAP v1.4 | Yu et al., 2017 | http://tanlab4generegulation.org/GMAP.html |
| Fit-Hi-C v2.0.7 | Ay et al., 2014 | https://github.com/ay-lab/fithic |
| RGT v0.12.1 | Gusmao et al., 2016 | https://www.regulatory-genomics.org/ |
| Python v2.7.15rc1 | / | https://www.python.org/ |
| R v3.4.4 | / | https://www.r-project.org/ |
| Yu et al., 2017 | Yu W et al. (2017) Identifying topologically associating domains and subdomains by Gaussian Mixture model And Proportion test. Nature Communications, 8, 535. |  |
| Ay et al., 2014 | Ay F et al. (2014) Statistical confidence estimation for Hi-C data reveals regulatory chromatin contacts. Genome Research. 24(6):999-1011. |  |
| Gusmao et al., 2016 | Gusmao EG et al. (2016) Analysis of computational footprinting methods for DNase sequencing experiments. Nat Methods. 13(4):303-9. |  |

